# *The biomedical sensor Cell-Fit-HD^4D^*, reveals individual tumor cell fate in response to microscopic ion deposition

**DOI:** 10.1101/2020.03.12.987347

**Authors:** M Niklas, J Schlegel, H Liew, DWM Walsh, F Zimmermann, O Dzyubachyk, T Holland-Letz, S Rahmanian, S Greilich, A Runz, J Debus, A Abdollahi

## Abstract

Here we present the biomedical sensor *<u>cell</u>-<u>f</u>luorescent <u>i</u>on <u>t</u>rack <u>h</u>ybrid <u>d</u>etector*^4D^ (*Cell-Fit-HD*^*4D*^) to reveal individual tumor cell fate in response to microscopic ion deposition in ion beam therapy. The sensor enables long-term monitoring of single tumor cells after clinical ion beam irradiation in combination with single-cell dosimetry. *Cell-Fit-HD*^*4D*^ is read out *in-situ* by conventional optical microscopy. Direct visualization of a clinical ion beam is hereby possible for the first time. The possibility to reveal fate of individual cells from a cell cohort demonstrates that our biomedical sensor clearly differs from conventional experiments that characterize cellular response after radiation on a population level. *Cell-Fit-HD*^*4D*^ is therefore used to mimics the clinical situation of a defined tumor depth during tumor treatment by ion beam therapy. Our biomedical sensor is able to provide crucial input for current mechanistic approaches to biophysical modelling of the effect of ionizing radiation on biological matter. In the clinical context, obtaining multi-dimensional physical and biological information on individual tumor cells is an important step to further transform ion beam therapy into a highly precise discipline within oncology.

## Introduction

Ion beam therapy with protons and heavier ions, such as helium, carbon and oxygen is transforming the field of radiotherapy into a highly precise discipline within oncology [1]. Highly localized energy deposition of an ion at the end of its trajectory, the so-called Bragg peak, enables precise deposition of dose inside a tumor, while sparing the surrounding healthy tissue in ion beam therapy [2, 3]. By combining multiple ion beams of coordinated energies, one is able to create a *spread out Bragg peak* (SOBP), an extended region of uniform dose in depth on a macroscopic scale. Heavy ions predominantly induce complex, persistent DNA Double-Strand-Breaks (DSBs), enhancing their relative biological effectiveness (RBE) in comparison to photon irradiation [3-5]. Despite the promising clinical success of ion beam therapy, there is still a great lack of understanding of the fundamental biological mechanisms that link the physical energy deposition by ions and the tumor response on molecular, subcellular and cellular scales.

Potential benefit of ions compared to conventionally used photons is often expressed by a single parameter, the RBE. The RBE is defined by the ratio of dose applied by photons and ions respectively to reach the same biological effect. On the cellular level, the dose applied in a SOBP is characterized by the entire spectrum of linear energy transfer (LET) and a Poisson-distributed number of actual ion traversals, as compared to the quasi-homogeneous photon dose [3]. As a consequence, cells in a given population do not only receive a distribution of “cellular” doses (i.e. dose per cell), but different cells receiving the same physical dose can be exposed to highly distinct number of ion traversals and LET combinations. Visualisation and spatiotemporal correlation of the energy deposition in a tumor cell, combined with a biological readout of its fate in a clinical ion beam, would increase overall understanding of the relationship between energy deposition and biological response.

Microbeams and CR-39 detectors have been used to study the effect of individual ion traversals on single cell fate [6, 7]. The microbeam experiments are not capable of fully elucidating the biological response of cells in a clinical ion beam field due to field’s complex energy spectrum as compared to a mono-energetic microbeam. To circumvent these limitations, the number of radiation-induced DNA damage foci (RIF) and their persistence in a cell nucleus have been used by others as a biological surrogate of the physical energy deposition in individual tumor cells [8]. The recording and analysis of RIFs present in a large cell population at a single time point in turn only represents a snapshot and thereby disregards the kinetics of DNA repair of individual cells triggered by the energy deposition in each individual cell.

The biomedical sensor *<u>cell</u>-<u>f</u>luorescent <u>i</u>on <u>t</u>rack <u>h</u>ybrid <u>d</u>etector*^4D^ (*Cell-Fit-HD*^*4D*^) enables long-term monitoring of single tumor cells after clinical ion beam irradiation in combination with single-cell dosimetry. The setup in which *Cell-Fit-HD*^*4D*^ is used mimics the clinical situation of a defined tumor depth during tumor treatment by ion beam therapy (Figure 1). The *Cell-FIT-HD*^*4D*^ sensor enables, for the first time, a complete deconvolution of the mechanisms linking the energy deposition of ions on single cell level with the ultimate and individual fate of heterogeneous tumor cells. In contrast to existing radiation experiments, the biomedical sensor combines single cell dosimetry with individual tracking of a large number of cells up to five days by long-term live cell imaging using widefield microscopy. This experimental approach enables investigation of fundamental mechanisms linking the microscopic heterogeneous energy deposition to the response of individual tumor cells. The biomedical sensor *Cell-Fit-HD*^*4D*^ will enable deeper comprehension of response of tumor irradiation by ion beams in the clinical context.

**Figure 1.**
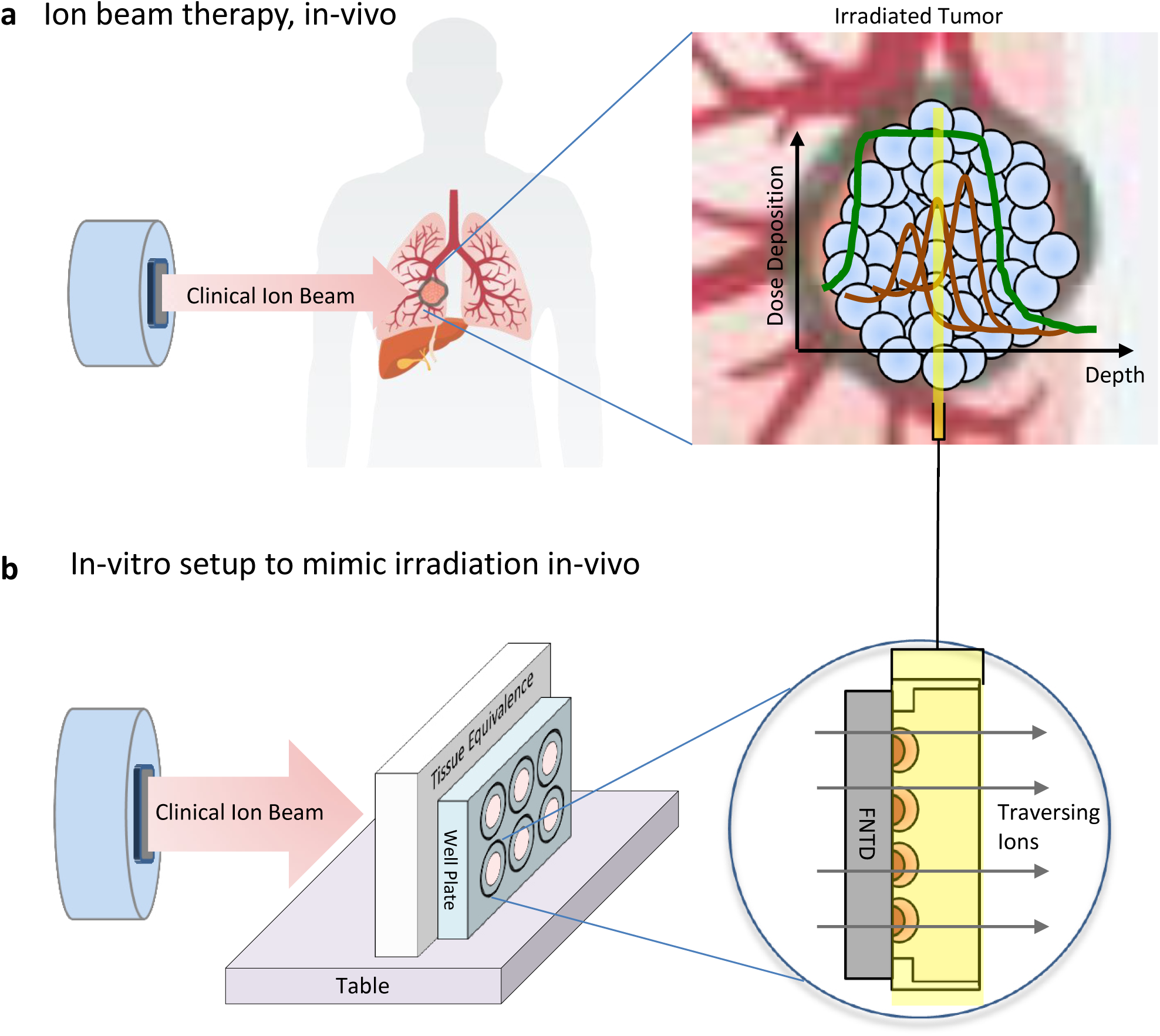
The biomedical sensor *Cell-Fit-HD*^*4D*^ is used to mimics the clinical situation of a defined tumor depth during tumor treatment by ion beam therapy. **(a)** Ion beam therapy is able to deposit the prescribed dose precisely in the tumor volume while sparing healthy surrounding tissue. By complex superposition of single Bragg peaks (ochre lines) a so-called spread-out Bragg peak (SOBP, green line) is formed. The SOBP enables a homogenous dose deposition in a small tumor volume element. On a microscopic level, every tumor cell localized in a given plane perpendicular to the beam (depicted by the yellow bar) is exposed to individual energy and hence dose deposition. **(b)** The design and irradiation setup of our in-vitro biomedical *Cell-Fit-HD*^*4D*^ mimics the clinical irradiation scenario of tumor cells in such a given plane (yellow bar). The fluorescent nuclear track detector (FNTD) in a shape of a thin wafer, is replacing the glass bottom of an ordinary multiwell imaging plate. A viable cell layer is seeded on the FNTD. Standard cell culture treatment is hereby possible. The incident ion beam has to penetrate stopping material in front of the multiwell plate, acting as healthy tissue equivalence in front of the tumor. Using this setup, the tumor cells of *Cell-Fit-HD*^*4D*^ are placed in the SOBP of the incident ion beam reflecting the clinical situation. Irradiation setup was used according to Dokic, Mairani et al. 2016.

## Materials and Methods

### Al_2_O_3_:C,Mg based FNTD

The fluorescent nuclear track detector (FNTD) was in the shape of a wafer of thickness 100 µm and of diameter 20 mm. The FNTD is a detector for single ion track visualization with sub-µm resolution [9, 10]. A detailed description of the FNTD can be found in [10, 11]. The FNTD is made of Al_2_O_3_:C,Mg. The crystal structure exhibit F^2+^_2_(2Mg)-centers fluorescent colour centres. They undergo transformation when capturing secondary electrons released by traversing ions. This leads to a unique foot print of the traversing ion in the FNTD. The color centers can be excited at 620 nm, prompting fast fluorescence at 750 nm. A read-out by confocal laser scanning microscopy enables visualization of individual ion traversals with sub-µm resolution. The FNTD is sensitive for ions with LET > 0.5 keV/µm [12, 13]. The detection efficiency is close to 100% for the ion spectra found in ion beam therapy [13].

### Irradiation of Cell-Fit-HD^4D^ in clinical setup

Irradiation of *Cell-Fit-HD*^*4D*^ was performed at Heidelberg Ion Beam Therapy Center (HIT) at Heidelberg University Hospital, Germany. The biomedical sensor was aligned perpendicular to the incident carbon ion beam (Figure 1b). The Irradiation was similar to [14]. The cell layer was positioned in the middle of the 1 cm wide spread-out Bragg peak (SOBP). The water equivalent depth was approximately 3.5 cm. Planned physical doses were 1Gy and 0.5 Gy, respectively. Both irradiations were principally identical except for rescaling of the ion beam fluence to gain defined physical dose. PMMA (thickness= 3 cm) was used as blocking material in front of the biomedical sensor. Tumor cells in the wells 1 and 4 (Figure 2a) were facing the ion beam. For the control additional PMMA block (thickness= 21 cm) was placed in front of the wells 3 and 6.

**Figure 2.**
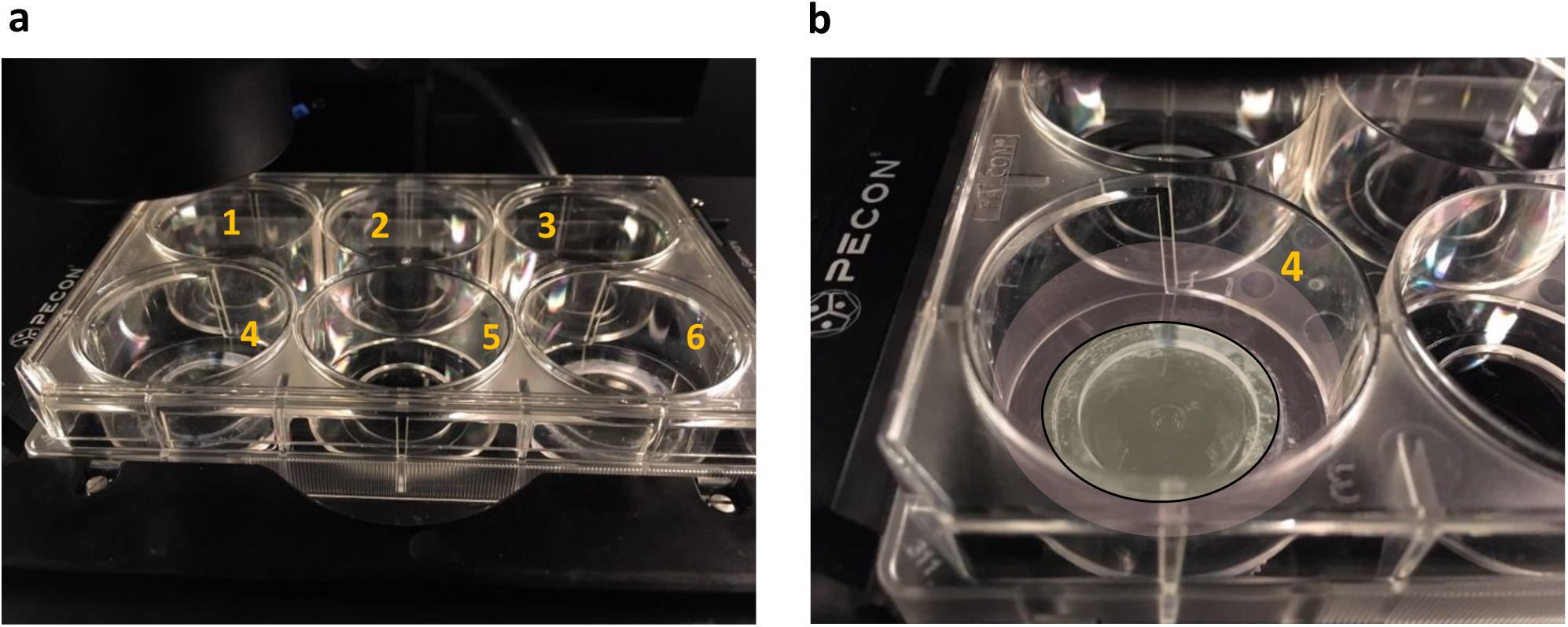
The design of the biomedical sensor *Cell-Fit-HD*^*4D*^ allows for live-cell imaging with optimal cell physiological condition. **(a**) The sensor consists of a conventional imaging plate. The well-bottom is substituted by a fluorescent nuclear track detector (FNTD). **(b)** The FNTD (indicated by the grey-colored disc) acts as device for detection of single ion traversals and as substrate for viable cell coating. In this exemplary experimental setup the bottom of well 4 comprises the FNTD (grey disc). The cell culture medium is indicated in magenta. The imaging light beam passes the FNTD from below.

### Read-out of Cell-Fit-HD^4D^

Except for the initial read-out, the physical compartment (FNTD) and the biological compartment (cell layer) are recorded independently.

The physical compartment was imaged by confocal laser scanning microscopy (CLSM, LSM710 ConfoCor3, Carl-Zeiss AG). A detailed read-out protocol is presented in [10, 15]. Briefly, tile scans consisting of overlapping imaging stacks were recorded: 40x oil objective, zoom= 1.1, number of rescans (line sum) = 2, dwell time= 6.3 µs. An imaging stack consists of 21 imaging planes of dimensions 1024 × 124 pixels (1 pixel= 0.189 × 0.189 um^2^) separated by 5 µm (without correction for refractive index mismatch) in axial (z) direction [16]. Transmission photomultiplier tubes (T-PMTs) were used to record the *spinels* in the transmitted-light channel; avalanche photo diodes (APDs) with long-pass filter (detection window > 650 nm) was used to record the physical track information (i.e. track spots) in the fluorescence channel in parallel. The spinels are crystal defects in the FNTD employed as landmarks for subsequent image registration (see *paragraph Stitching and registration procedure below*). Imaging started 5 µm below the FNTD surface.

The biological compartment was recorded by inverted widefield microscopy (IX83, Olympus), including 20x/NA 0.8 air objective, illumination system Lumencore spectra X LED system, Hamamatsu Orca flash V2 sCMOS camera (exposure time = 4 ms and 200 ms for brightfield and fluorescence channel, respectively), filter-set, and an incubation chamber (humidified atmosphere, standard culture conditions: 37 °C, 5% CO2). Image stacks at several positions were recorded. A stacks contained 3 planes of dimensions 2048 × 2048 pixels (665.6 × 665.6 μm^2^, pixel size of 0.325 × 0.325 μm^2^) separated by 1.5 µm (without correction for refractive index mismatch). The vertical position of the cell layer was detected by an autofocus routine.

In the initial read-out (Figure 3) the physical and biological compartment were scanned sequentially in a single step by widefield microscopy to record the *spinels* (FNTD) and biological information (i.e. cell layer). The scan comprised imaging stacks of several positions. Each stack were covering the total FNTD and the cell layer in vertical dimension of *Cell-Fit-HD*^*4D*^) of dimensions 2048 × 2048 pixels (665.6 × 665.6 μm^2^, pixel size of 0.325 × 0.325 μm^2^) separated in z by 5 µm (without correction for refractive index mismatch).

**Figure 3.**
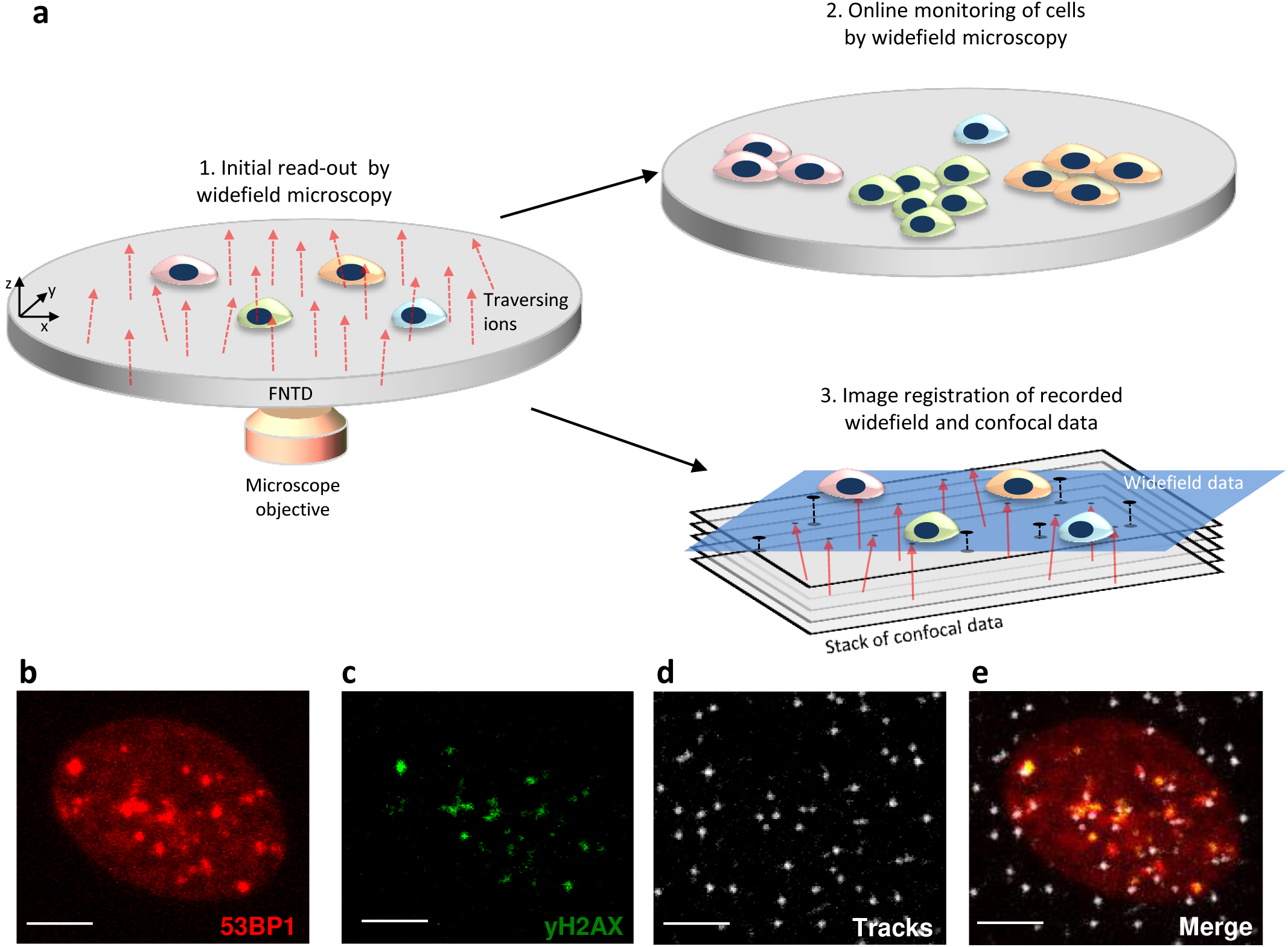
The workflow of *Cell-Fit-HD*^*4D*^ enables online-monitoring of individual cellular response to ion irradiation and a subsequent correlation to the physical energy deposition on sub-um scale. **(a)** Workflow: Immediately after ion irradiation, *Cell-Fit-HD*^*4D*^ is transferred to the widefield microscope. (1.) By initial read-out, the FNTD and the tumor cells are scanned in a single step. Thus the earliest response of the tumor cells to irradiation (fluorescent signal) as well as the crystal defects (spinels) in the wafer (transmitted light) acting as landmarks for later image registration are recorded. (2.) In a second step the online monitoring of the biological compartment is performed up to five days by widefield microscopy. Typical acquisition intervals are 45 min to probe the cellular and molecular dynamics. (3.) After online-monitoring, the *Cell-Fit-HD*^*4D*^ is transferred to a confocal laser scanning microscope to read out the FNTD. Using the recorded fluorescent track spots (i.e. unique foot prints of the traversing ions left in the FNTD) and image registration routines (crystal defects are indicated by black discs), each ion track (red arrows) can be reconstructed into the cell layer recorded in the initial read-out with sub-µm precision. **(b)** Fluorescent signal of 53BP1 damage protein in A549 cell nucleus recorded in the initial read-out approx. 5 min after carbon ion irradiation with planned physical dose of 1.5 Gy. The bright spots (termed RIFs) are indicating accumulation of 53BP1 at the DNA damage sites. **(c)** Immunfluorescent staining of yH2AX confirms specificity of 53BP1 RIFs. **(d)** Fluorescent read-out signal of the FNTD. Each bright track spot correspond to a single ion track. **(e)** Spatial correlation of fluorescent 53BP1 and yH2AX signal, and the ion tracks in a cell nucleus. Scale bars, 5 µm.

Concerning the read-out of the Mounting-architecture, the FNTD was mounted in a glass bottom dish (MatTek Corporation, 6 well, Part No. P06G-1.5-20-F) and was recorded as described above.

### Cell culture

A549 human Non-Small-Cell-Lung-Carcinoma (NSCLC) cells (ATTC^®^ CCL-185) were cultured using Dulbecco’s Modified Eagle’s Medium (DMEM) supplied with 10% Fetal Bovine Serum (FBS) and 1% Penicillin/Streptomycin at 37°C, 5% CO_2_. For live cell imaging, respective medium lacking phenol-red was used. A549 cells were retrovirally transduced with a construct coding for the N-terminus of 53BP1 fused to the sequence coding for fluorescent mCherry-protein (Addgene Catalog # 19835; originally described in [17] and selected with puromycin (5µg/ml;). Puromycin-selection was applied on a regular base during culturing as well as during the experimental course.

### Stitching and registration procedure

A detailed description of the stitching and registration procedure can be found in [15]. A single imaging field recorded by the widefield microscope (20x objective) comprised 4×4 tiles recorded by confocal microscopy (40x objective, zoom= 1.1). The zoom was set to 1.1 to decrease vignetting effects at the imaging margin. For image processing purposes the single tiles comprising an imaging stack of 20 layers were normalized using the in-house written software called *FNTD package* [18]. All tiles were recorded with a spatial overlap of approximately 20 × 193.5 µm^2^. A binary mask was computed comprising the segmented track spots as foreground objects of the maximum intensity projection of each imaging stack (using 16 imaging planes). To create a binary mask a sequence of thresholding (threshold value of 0.2 of the normalized data), filling of regions and holes and replacement of small objects (<= 4 pixel) was applied to the raw data. The segmented track spots are acting as unique fingerprints for the subsequent stitching process. In a first step all neighboring tiles of the 4×4 scan were stitched. Using the stitching parameters a complete row (comprising 4 tiles) were reconstructed. Each row was cropped at its upper imaging margin (approximately 9 µm) to erase areas with zero intensity introduced by the stitching. In a second step all neighboring rows were stitched. Finally the imaging field was reconstructed using the stitching parameters of the neighboring rows. The accuracy of stitching is directly reflected in the subsequent image registration process. For additional quality control an overlap of the stitched tiles was visualized in red green color coding. The spatial overlap was thus visualized in yellow.

For the registration of the confocal and widefield imaging data the *spinels* recorded in the transmission light channel were used as landmarks. The intensity-weighted centroid of each *spinel* was computed in the maximum intensity projection of in the confocal (input) and widefield (base) imaging stacks which were initially stitched. For the registration process a *projective transformation* was used. All *spinels* detected were used as landmarks. To control the accuracy of transformation the Euclidean distance between the *spinels* in the input and base image acting as control point pairs (*cpps*) as well as the four corner angles of the registered confocal imaging field were calculated. The mean distance between cpps was on average less than 0.4 µm for all imaging fields registered. The corner angles of the registered imaging fields were in the range of 90° +-0.3°.

### Ion track reconstruction and dose calculation in cell nucleus

#### 1. Ion track reconstruction in FNTD and extrapolation into cell layer

The images obtained from the FNTD were analysed by in-house written software called *FNTD package* [18]. This software can be run as a plugin for the image-processing and analysis software *ImageJ/Fiji*. The software is capable of identifying track spots from ions. Based on user input the software can link these track spots together to reconstruct the ion track in the FTND. For each series of images obtained, the parameters specified in the *FNTD package* were chosen iteratively in order to maximize the number of tracks found visible on the image stack while minimizing the number of false tracks arising from linking background noise and delta electrons. After the track is reconstructed in 3D in the FNTD, the trajectory was fitted into a line in the 3D space using linear regression in the software *R [19]* and the slope of the trajectory was obtained both in xz and yz directions with error intervals. These slopes were then used to extrapolate the ion tracks into the cell layer at vertical position of 2 µm (thickness of cell layer ∼ 4 µm) with respect to the FNTD surface.

#### 2. LET and dose calculation

The average intensity of each track was obtained from the *FNTD package*. Distribution of the corresponding track intensities *I* show two distinct peaks and they are related to LET of the ion *j* by

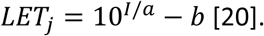

A Monte Carlo simulation of the ion spectra at the FNTD surface was performed in the FLUKA Monte Carlo code. The parameter *a* and *b* were chosen by matching the peaks of the primary (C-12) ion with the Monte Carlo results, while optimizing for total dose by C-12 ions, total number of C-12 ions and LET distribution of C-12 ions. After the LET of individual ion track was calculated, the dose of each nucleus (*dose*_*i*_) was calculated according to

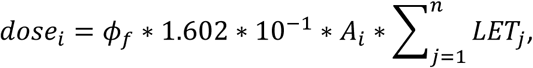

with *A*_*i*_ being the cross-sectional area of *nucleus*_*i*_, and *ϕ*_*f*_ being the fluence factor. The fluence factor is defined as the fluence of each individual track in the FNTD (as obtained from the FNTD package) multiplied by the area scaling factor (area of the FNTD divided by area of imaging field of the cells), in order to account for the fact that a different area is used to calculate the fluence in the cell layer. To determine the ion traversals *j =* 1 … *n* for each cell nucleus individually the intersections of the track endpoints with the segmented cross sectional areas *A*_*i*_ (see below) were calculated. LET and dose calculation was done separately for 1 Gy and 0.5 Gy irradiation. Here, we focused on the dose deposition by primary (carbon ions, C-12) ions. Approximately 94 % of the energy is deposited by the primary ions [21]. Based on the histogram of the intensity images converted to LET spectrum the primary and secondary ions were distinguished from another by introducing an LET threshold (LET_min_). Expected (from the Monte Carlo (FLUKA) simulation [14]) and reconstructed values are listed in Table 1. We separated the total dose in the entire imaging field defined by the widefield microscopy (FOV) from the reconstructed dose values in the cell nuclei.

**Table 1:** Comparison of beam parameters: expected by Monte Carlo simulation and experimental ones.

| Irradiation | 0.5 Gy, C-12, SOBP |  | 1 Gy, C-12, SOBP |  |
| --- | --- | --- | --- | --- |
|  | Expected | Experiment | Expected | Experiment |
| LET<sub>min</sub> for C-12 ions [keV/μm] | ~ 44.50 | 43.90 | ~ 44.50 | 44.42 |
| FOV: **Total dose** by C-12 [Gy] | 0.47 | 0.41 | 0.94 | 0.87 |
| FOV: **Fluence** of C-12 ions [cm<sup>-2</sup>] | 4.16 e+06 | 4.14 | 8.33 e+06 | 8.43 e+06 |
| Cell Nuclei: **median dose** by C-12 [Gy] | -- | 0.45 | -- | 0.90 |

### Post-time lapse imaging staining

Following live-cell microscopy, culture medium was immediately aspirated and cells were washed with PBS, followed by fixation with ice-cold 70% EtOH and storage at −20°C for a minimum of 24h. For protein staining, cells were washed with PBS, and concomitant blocking and cell permeabilization was conducted with 3% BSA, 0.2% TritonX-100 in PBS for 20 min. Primary antibodies (mouse anti-yH2AX, Cell Biolabs, STA-321, 1:100; rabbit anti-53BP1, Cell Signaling Technology, 4937S, 1:200; mouse anti-p21, Santa-Cruz, sc-6246, 1:50) were diluted in washing buffer (0.6% BSA, 0.02% TritonX-100 in PBS) and applied on top of FNTD for overnight incubation at 4°C. Wells were consecutively washed two times for 5 min in washing buffer before incubation with secondary antibodies (Alexa Fluor-488 goat anti-mouse IgG, Invitrogen, A-11029; Alexa Fluor-647 goat anti-mouse IgG, Invitrogen, A-21236, Alexa Fluor-488 donkey anti-rabbit IgG, Invitrogen, A-21206; all 1:400) for 5 h at 4°C. Cells were washed sequentially with washing buffer and PBS for 5 min each and stored in fresh PBS at 4°C until microscopic acquisition.

### Cell segmentation and tracking

Cell segmentation and tracking are performed using a hybrid algorithm that combines features of both tracking-by-detection and model-based tracking approaches. Cells on the first image of the stack are segmented using the segmentation approach described in [22]. Briefly, this algorithm consists of three core steps: 1) Initial (non-PDE-based) segmentation of the foreground; 2) Splitting the foreground into separate instances (cells); and 3) Refinement of the segmentation by multi-level-set framework [22]. Segmentation of each consecutive image in the stack follows the same logic. However, implementation of the first two steps is different as it relies on the information from the previous time point. Namely, the foreground mask (step 1) is estimated using the threshold that best preserves the foreground region obtained as the final result of the segmentation of the previous image in the stack. An analytic expression for this threshold can easily be derived using the intensity histogram of the image. Next, for splitting the foreground into separate regions (step 2), we also use the calculated segmentations of the cells from the previous time point. The foreground is first split into super-pixels, calculated using elliptic features, resulting in an over-segmented image. The super-pixels are consecutively assigned to one of the cells depending on the amount of the overlap between them and regions occupied by each of the cells. In this, detected regions that were not associated with any of the existing cells are labelled as “new” cells and the ones that were not associated with any of the super-pixels are labelled as “disappeared”. This process results in a rough segmentation that is subsequently refined using the multi-level-set framework [22](step 3).

### Data processing

If not declared otherwise the data processing was performed using *Matlab* (R2019a).

## Results

Former versions of *Cell-Fit-HD* were published in [14, 23, 24]. Here, we aimed at observation and correlation of the long-term cellular response (up to five days) of single tumor cells to their individual energy exposition with sub-µm resolution after clinical ion beam irradiation. For that purpose, we developed a technique that combines widefield time-lapse microscopy with ion track reconstruction from FNTD, the results of which we present in the following as the biomedical sensor *Cell-Fit-HD*^*4D*^.

### Methodological development and validation

#### Design of Cell-Fit-HD^4D^ and manufacturing approaches

For the current design of our biomedical sensor *Cell-Fit-HD*^*4D*^ we substituted the glass bottom of a conventional 6-well imaging plate with a fluorescent nuclear track detector (FNTD) in the shape of a wafer and seeded cells on top, as for usual cell culture experiments. In this design of the hybrid detector the cell layer (biological compartment) is directly adherent to the FNTD (physical compartment, Figure 2).

For the manufacturing approach, we cleaned the surfaces of the imaging plate with ethanol (70%) to disinfect them and to prevent interfering particles. Small droplets of histo-acryl glue (B. Braun GmbH, Art. Nr. 9381104) - in total 8 µl - were placed on the circular-shaped frame of the bottom of the well bottom. We found that this applied volume minimizes the chance of contamination of the inner well with histo-acryl, while ensuring solid and enduring stability. The sterilized wafer (FNTD) was gently pressed by means of a suction device onto this circular frame (adhesive surface). The number and position of the wells to be modified by FNTD wafer depends on the experimental setup. In Figure 2, the bottom of well 4 and well 6 were replaced by a FNTD wafer respectively. Well 4 contained the irradiation cohort and well 6 acted as control. We found that upside-down placement of the modified imaging plate in a conventional cell culture incubator (37°C; 5% CO_2_, humidity) for approximately one minute is sufficient to allow complete polymerization of histo-acryl. Since potential glue residues could occasionally be detected, both surfaces of the wafer were cleaned by help of a cotton bud ear stick soaked in ethanol (70%). This further reduced the probability of later detection of microscopic glue residues in the cell medium. Quality control of the cleaned wafer surfaces was performed by conventional bright-field microscopy (magnification: 20x). As we found that extensive well rinsing further improves cell viability, we additionally sterilized and washed the modified wells with 1ml ethanol, followed by 2x 1ml PBS. Robust fluid retention was confirmed by incubation of the multiwell plate filled with cell medium in the incubator for approximately 30 min. We noticed that this test incubation needed to occur at experimental conditions (37°C; 5% CO_2_, humidity), as temperature and humidity could influence glue stability. A549 cells were then seeded on top of wafer forming the biological compartment. Afterwards, the plate was placed in the cell culture incubator overnight (o/n). The general biocompatibility of the FNTD to allow for adherent cell layers was demonstrated in [23, 25].

In contrast to earlier designs of *Cell-Fit-HD*^*4D*^, the imaging light beam by optical microscopy here passes through the wafer from below for cell acquisition (Figure 2b). We found that this leads to optimal cell physiological conditions in terms of oxygen-, nutrient-, and spatial supply, which was tested by recording of proliferation curves over the course of days. The utilization of a widefield microscopy allows for fast acquisition of hundreds of cells while minimizing phototoxicity and therefore maximizes cell viability and observability of the radiation responses. Additionally, the new design completely abolishes the need for any kind of handling between irradiation and acquisition, since the biomedical sensor can be transferred directly to the microscope after irradiation.

### Workflow of Cell-Fit-HD^4D^

Approximately 18 h before irradiation A549 cells were seeded on top of the wafer using DMEM. We experimentally determined 150,000 cells/well to be a good seeding number for sufficient number of cells per imaging field while allowing for population expansion over the course of several days. We further noticed that, for optimally uniform cell distribution, incubation at room temperature for approximately 20 min is helpful to allow cells to settle down without influence of hydro-dynamics. The biomedical sensor was then stored in the incubator o/n to allow for the adherence of a cell layer. Before irradiation cells were monitored by microscopy to ensure attachment on the modified well bottom and normal physiological morphology. Unusual morphology or floating cells were a sign for contact of glue residues with the medium. During method development, we started to replace the conventional DMEM cell medium with pre-warmed phenol-free DMEM prior to irradiation, as this would decrease background signals during microscopy.

The tumor cells were placed in the SOBP of carbon ion (C-12) irradiation field. The biomedical sensor (including its cell layer) was placed perpendicular to the incident ion beam (Figure 1b). To ensure no leakage of the culture medium during irradiation, we found out that it was usefuel to seal the wells with Parafilm (Pechiney Plastic Packaging Inc., USA, Cat. No. PM996) under slight pressure applied by a sheet of a wiping paper under the plate lid. This construction was maintained stable by additional wrapping of the multiwell plate sides with Parafilm. In the presented case, well 6 (Figure 2a) acting as control group was blocked from the incoming ion beam by placing a PMMA block of 21 cm thickness in front of the multiwell plate. Immediately after ion irradiation, *Cell-Fit-HD*^*4D*^ was transferred to the temperature controlled (37°C, 5% CO_2_) widefield microscope. During this step the Parafilm was removed.

To guarantee a minimum delay between irradiation and acquisition start, all parameters for the microscope read-out (using *ScanR, Olympus*) were set in advance. In an initial read-out, the FNTD and the tumor cells were scanned in a single step (step 1, Figure 3a). Thus the earliest response and the initial position of the tumor cells to irradiation (fluorescent signal) as well as the *spinels* in the wafer (transmitted light) acting as landmarks for later image registration were recorded (*Materials and Methods, Stitching and registration procedure*). The initial read-out is crucial to enable precise spatial correlation of physical energy deposition and cellular localisation. (*Supplementary Information, Precision of microscopy stage movement*). In a second step the live cell imaging of the biological compartment was performed up to five days by widefield microscopy (step 2, Figure 3a-c). Typical acquisition intervals were 45 min to probe the cellular and molecular dynamics. A sequence of shorter time intervals of 15 min – in total 2 h - was added at the start of the imaging process to probe the early dynamics of 53BP1 after irradiation. Cells were still viable after five days of live-cell imaging. We found out that cell tracking is greatly impaired by high cell density at later observation time points (*Supplementary Information, Tracking using LSetCellTracker).* While cells were in principle still viable after five days of live-cell imaging, we found that cell tracking is greatly impaired by high cell density at later observation periods (*Supplementary Information, Tracking using LSetCellTracker*).

After live cell monitoring (i.e. at the time point of interest), the *Cell-Fit-HD*^*4D*^ was transferred to a confocal laser scanning microscope (CLSM) to read out the FNTD (step 3, Figure 3a,d). We could confirm that the FNTD can be scanned for ion track reconstruction without destroying the cell layer which was fixed after the online monitoring (ethanol 70%). The decoupling of live cell monitoring by widefield microscopy and ion trajectory reconstruction by recording the FNTD by CLSM presents the key achievement in this workflow. Spatial correlation of single tumor cells with the reconstructed ion tracks penetrating the tumor cells was then feasible by image registration of the read-out signals from the FNTD and cell layer recorded during the initial read-out (Figure 3e) [15]. Potential error sources – e.g. by cell migration directly after irradiation – in the determination of the physical energy deposition in individual tumor cell was minimized by a very short time interval of approximately 5 minutes between irradiation and live cell monitoring [15]. The landmarks (*spinels*) for image registration were recorded in the initial read-out directly after irradiation by widefield microscopy, as well as during the read-out of the FNTD by CLSM which was conducted after the time-lapse run. Using the recorded fluorescent track spots (i.e. unique foot prints of the traversing ions left in the FNTD) and image registration routines (*spinels* are indicated by black discs in Figure 3a), each ion track (red arrows) could be reconstructed into the cell layer recorded in the initial read-out with sub-µm precision [15, 16] (see also *Supplementary Information, Assessment of error sources/ uncertainties in spatial correlation*). Based on the intensity of a track spot the corresponding LET of the traversing ion in the FNTD and finally the dose deposited in the cell nuclei was calculated (Figure 4).

**Figure 4.**
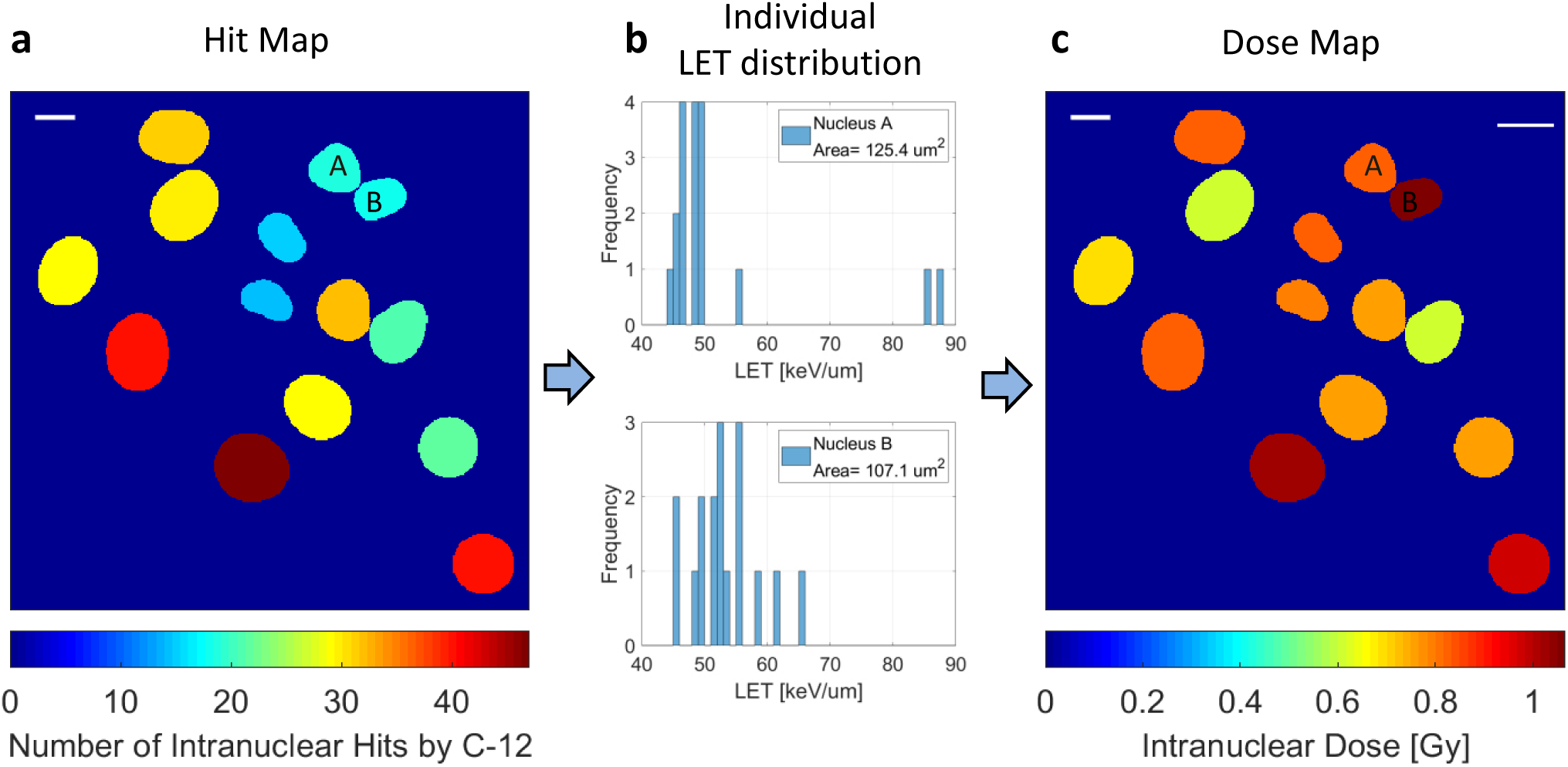
*Cell-Fit-HD*^*4D*^ allows to resolve the inhomogeneous energy deposition of clinical ion fields on the cellular level. **(a)** The cross-sectional areas of exemplary cell nuclei derived from the recorded 53BP1 signal are shown. The number of intranuclear ion traversals is encoded by the color of the cell nucleus. The computation of the nucleus area was conduced by *LSetTracker*. **(b)** The corresponding linear energy transfer (LET) distributions by the carbon ion traversals in nucleus A and B are depicted. **(c)** Based on the LET distribution for each nucleus and its cross-sectional area the physical dose is assessed for each nucleus. As shown here, despite similar hit numbers in A and B the corresponding dose level in the nuclei differ due to their individual LET distribution. We focused on the primary particles (i.e. carbon particles, C-12) depositing ∼94 % energy of the incoming ion beam. Scale bars, 10 um.

The RIFs (53BP1 foci) emerging after irradiation were segmented by *Trainable Weka segmentation* plugin for *ImageJ* [26]. Tracking and segmentation of individual cells recorded in the live-cell imaging were performed using *LSetCellTracker [22]*. After completion of the experiment, the biomedical sensor could be dissembled and the FNTD could be reused after bleaching.

### Preliminary biological results

#### Linear correlation of 53BP1 foci and deposited energy

Using of our biomedical sensor system, we investigated the correlation of the ions’ intranuclear energy deposition and the molecular response in form of DNA damage repair dynamics in individual A549 tumor cells.

The physical dose deposited in single cells is mainly determined by two variables: The number of traversed ions (especially primary C-12 ion hits accounting for ∼94% of the dose) and second, the sum of their specifically transferred energy within the nucleus, determined by the ions’ LET (Figure 5, SI5). We expected that due to the spectrum of LET present in the SOBP as well as substantial differences in nuclei sizes, the correlation between the number of ion traversals and the dose deposited in single cells can substantially deviate from linearity (Figure 5d). We therefore proposed a novel estimate ΣLET, i.e. sum of all ions’ LET traversing cell nuclei, to account for the broad LET spectrum of the traversing ions (Figure SI5a,b). This parameter disregards the nucleus area (unlike dose) and therefore accounts for the single cell as the most basic biological, integer unit.

**Figure 5.**
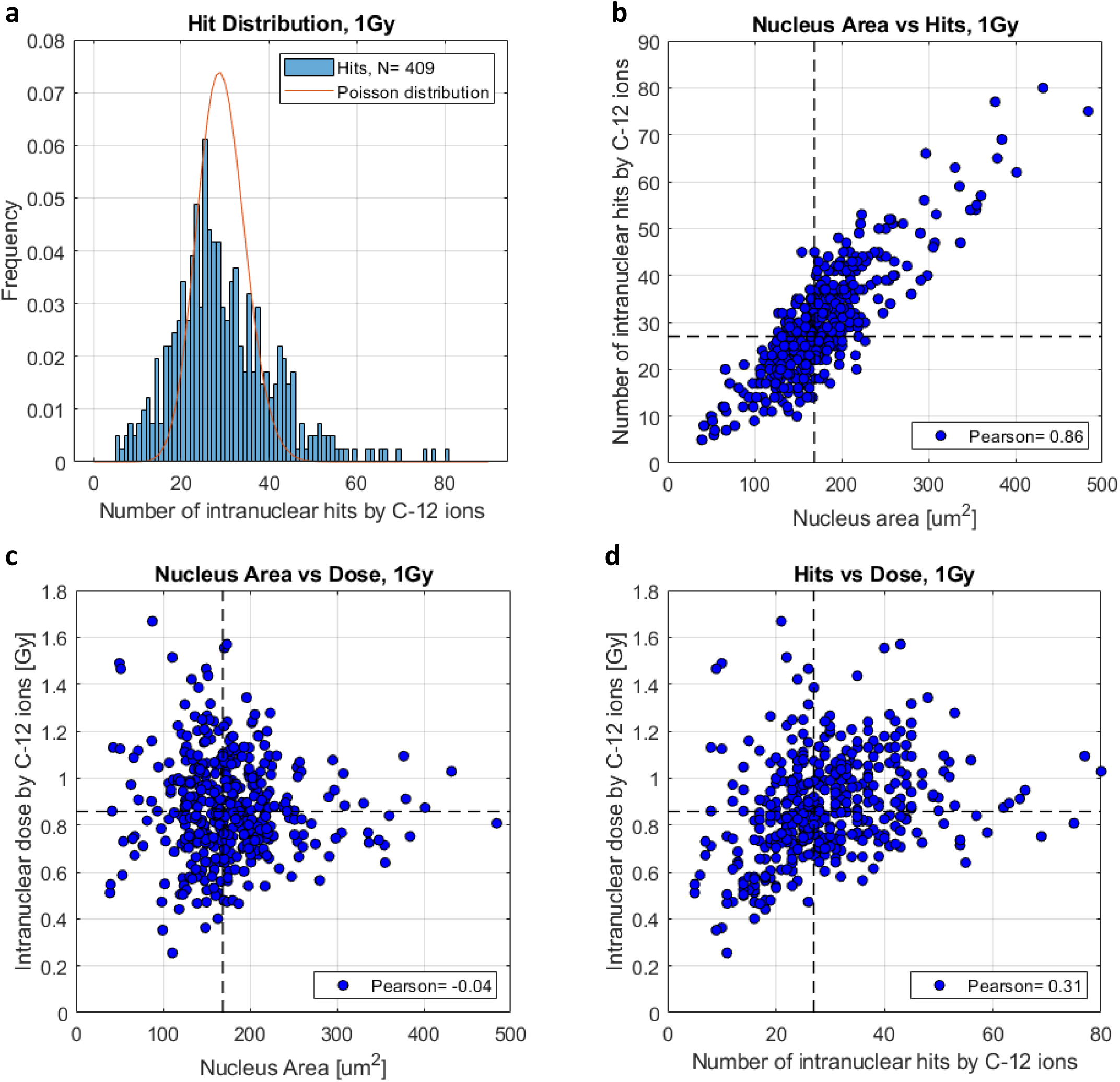
*Cell-Fit-HD*^*4D*^ enables to resolve the distribution of the physical beam parameters – hits and dose – on the cellular level. The correlation of both parameters can substantially deviate from linearity. **(a)** Distribution of number of intranuclear ion hits by carbon ions (C-12, 1Gy SOBP). The ochre dotted line indicates the corresponding Poisson distribution (λ = 29.2). **(b)** Correlation between the nucleus area (median area= 169 um^2^) and number of intranuclear ion hits (median hits= 27). **(c)** Correlation between the nucleus area and intranuclear dose (median dose= 0.86 Gy) deposit by the C-12 ions. **(d)** Correlation between the number of intranuclear ion hits and intranuclear dose. The Pearson-r-coefficients is stated for each correlation. The dashed lines indicate the corresponding median values and were used to define the quadrants.

The Poisson-distributed number of ion traversals (Figure 5a) in combination with the LET spectrum inside the SOBP (Figure 4, SI5) resulted in cell-specific combinations of primary ion hits and dose, where ion hit number did not necessarily translate to respective dose in a strictly linear manner. To elucidate the specific impact of each measure (number of ion traversals and dose received) we separately investigated the relationship of hits as well as dose to detectable 53BP1-RIF in single cells over time.

While we in principle observed positive correlation with RIF number for both, primary ion hits as well as dose, the data was characterized by substantial variability in the RIF-evocation for single cells which received equal dose or hit numbers at any given time (indicated by error bars in Figure 6b and SI6b). This might indicate major impact of the stochastic nature of energy deposition by ions on the cellular and sub-cellular level on RIF induction by heavy ions [27], as well as single cell variability in RIF formation caused by molecular or biological background heterogeneity of tumor cells. Remarkably, while the overall RIF number increased to a maximum within the first 1 h (Figure SI1), a one-to-one relationship of traversed carbon ions and elicited DNA damage foci formation could not be determined at any time point, indicating that not each single carbon hit provokes a complex DSB at the energies used in the presented experiment.

**Figure 6.**
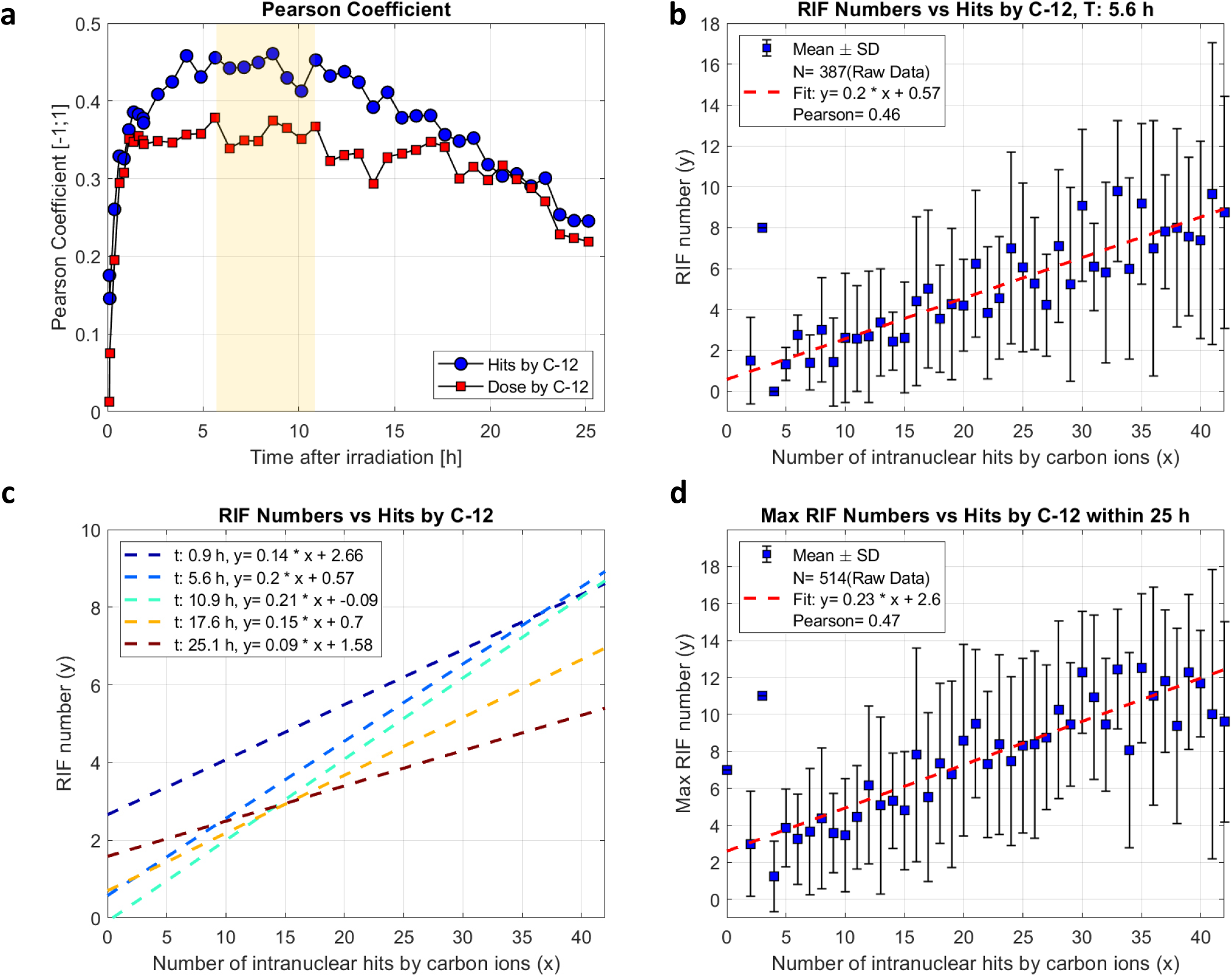
*Cell-Fit-HD*^*4D*^ enables correlation of physical ion beam parameters with molecular dynamics of individual tumor cells. The linear correlation between hits and RIFs peaks in the interval [5.6; 10.9] h post irradiation. This correlation is improved by calculating maximum number of RIFs in a nucleus within 25 post irradiation. **(a)** Time-dependent Pearson-r correlation coefficient within 25 h post irradiation. It indicates an initial fast increase followed by a plateau and subsequent slow decrease in correlation between number of intranuclear hits/ dose by carbon ions (C-12, planned physical dose of 0.5 and 1 Gy) and number of RIF (53BP foci) per nucleus. The maximum is in the interval [5.6; 10.9] h post irradiation (yellow bar). **(b)** Linear correlation between number of intranuclear hits by C-12 and the number of induced RIFs 5.6 h after irradiation. Statistically speaking five C-12 ion hits are needed to induce a single RIF. **(c)** Time-dependence (0.9, 5.6, 10.9, 17.6, 25 h post irradiation) of linear correlation between number of intranuclear hits by C-12 and the number of induced RIFs. **(d)** The maximum number of emerging RIF within 25 h after irradiation showed improved linear correlation compared to the number of intranuclear hits. The slope raised to 0.23 RIF/hit. For the computation of the Pearson-r correlation and linear regression analysis the raw data was used. Here, hit numbers were limited to maximum 42 hits in (**a-d**) to ensure sufficient number of data points. Only mother cells were considered in the data analysis.

To compare the RIF correlations with hits and dose, we calculated Pearson-r coefficients for the first 25h of observation and did linear-regression analysis. For both parameters, the coefficient was highest in the time interval between 5.6 h and 10.9 h post irradiation with values in the interval [0.42;0.46] for hit number and [0.29;0.33] for dose correlation (Figure 6a). Notably, the slope of the linear regression depicted similar progression over time as the Pearson’s-r coefficient with the same plateau for maximum slope values: [0.19; 0.21] RIF/hit and [5.13; 5.80] RIF/Gy (Figure SI6a). Hence, increasing variable-dependency appears to be accompanied by stabilization of the correlation. Hit-RIF-correlation provided higher Pearson-r coefficients compared to dose-RIF correlation. This was mainly attributed to greater fluctuations above 0.8 Gy in the dose-RIF-correlation (Figure SI6b). We therefore understand the number of ion hits to be a stronger predictive measure of complex DSB induction in these clinical ion fields, with about one RIF observed for every fifth carbon hit (disregarding its LET). We, therefore, focused in this manuscript on the parameter hit. Still, identical calculations were performed for the parameter dose (Figure SI6), although they will not be reported in this manuscript.

Individual RIF exhibit distinct, differential formation- and repair kinetics due to several reasons such as damage complexity, DNA accessibility. We therefore expected that a population level analysis at a single time point will never score maximum RIF numbers for all single cells, and thereby cannot indicate the true amount of induced complex damage. In contrast, utilizing our data from the implemented live cell microscopy, we were able to assess the maximum number of RIFs observed in each individual cell. Introducing maximum RIF numbers slightly improved the linear correlation to the number of hits (Pearson-r coefficient= 0.47), and elevated the induction rate to 0.23 RIF/hit (Figure 6d). This correlation therefore provided a better surrogate for the physical energy deposition compared to determination of RIF numbers at a single time point and differs from broadly applied measurements of RIF as irradiation surrogate 15 or 30 min post-irradiation. Notably, in regard to the above mentioned finding that linearity peaks in the interval between 5.6 h and 10.9 h post-irradiation, we found that only 74% of the recorded cells showed highest RIF numbers within that respective time span (Figure SI7a), pointing towards stabilization of linear dependence of RIF development in trailing cells. The raw data of maximum RIF number and additional correlation to the parameter ΣLET are depicted in Figure SI7.

### Clonogenic potential and growth arrest induction

Assessment of dose-dependent clonogenic survival of irradiated cells presents the gold standard for evaluation of radiation sensitivity. However, it is not clear how microscopic energy deposition (i.e. on a cell nucleus scale, Figure 4) in a clinical ion beam field links to single cell fate determination. We therefore semi-automatically tracked cells over a time frame of > 4 days and correlated their clonogenicity (total number of progeny cells from individual mother cell) to individually received primary hit numbers (Figure 7a,b). While a certain tendency towards negative impact of hit number on proliferation could be suspected, the data did not depict that ion hit number directly translates to proliferative arrest in a quantitative manner (Pearson-r coefficient= −0.21, Figure 7b). This is a hint for additional processes on the biological level which comes into play in the induction of clonogenic death of tumor cells by heavy ions. Clonogenic death of cells does not solely reflect microscopic hit (Poisson-) distribution (Figure 5a).

**Figure 7.**
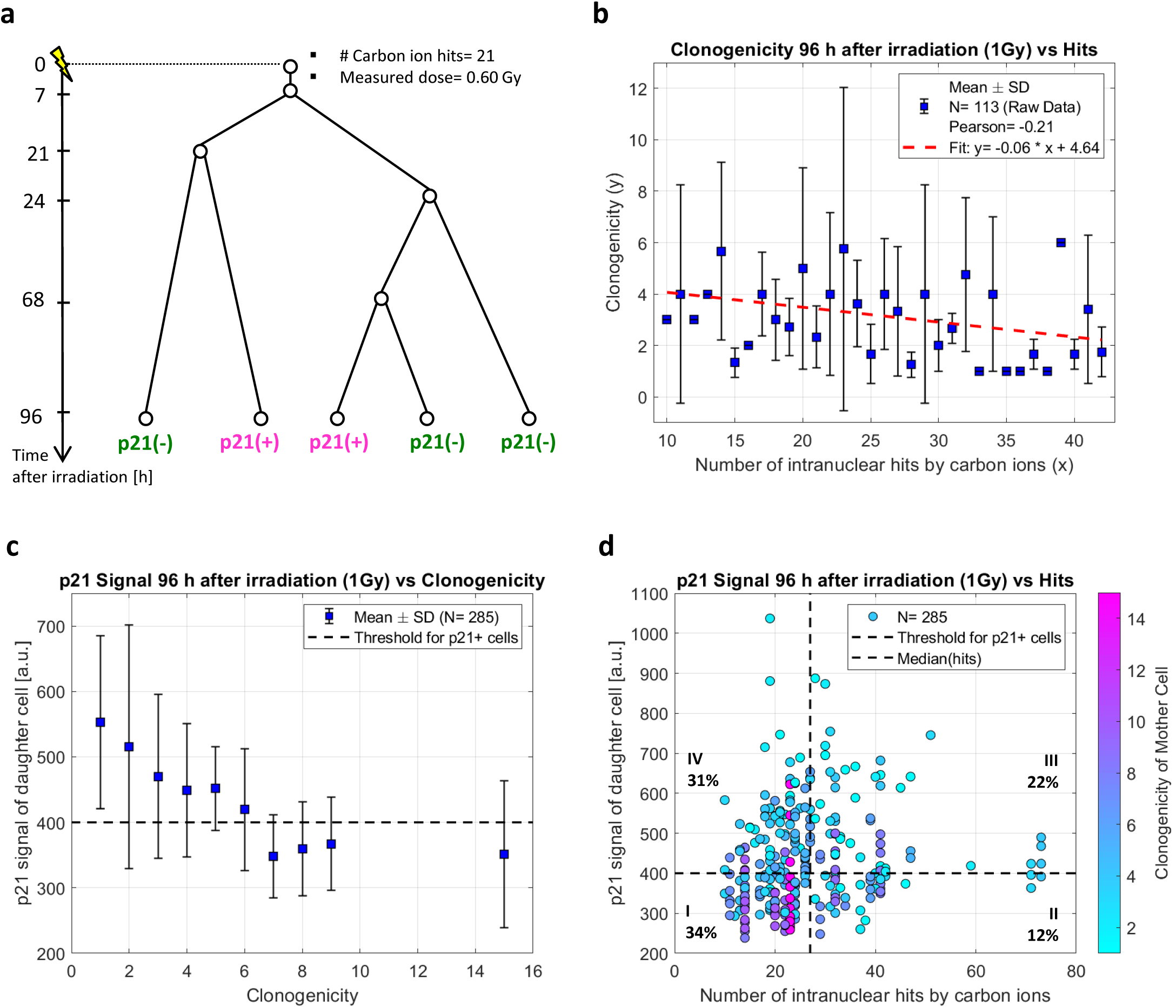
*Cell-Fit-HD*^*4D*^ combines single-cell dosimetry with molecular and cellular dynamics of individual tumor cells. **(a)** Exemplary cell division tree of an irradiated cell (by carbon ions, planned physical dose of 1 Gy). Irradiation took place at time point zero. P21 positive cells are highlighted in magenta; p21 negative cells in green. **(b)** Correlation of clonogenicity (number of progeny cells deriving from a single mother cell) with the number of intranuclear carbon ion hits. A tendency in the decline of proliferation with increasing hit numbers is visible. For linear regression analysis raw data was used. **(c)** Correlation of p21 signal in cell nuclei with clonogenicity. A binary distribution with thresholding of 400 a.u. is mirroring the binary read out of p21-positivity and -negativity. **(d)** By multi-dimensional correlations we were able to define combinations of physical beam (hits)- and molecular (p21 status 96 h after irradiation) parameters to predict proliferation potential and to find potential marker for radiation-resistance. The plot was divided into four quadrants (I-IV): median hits= 27, p21 threshold= 400 a.u. The portion of the total population is given for each quadrant.

Our biomedical sensor allows for fluorescent staining of the fixated cell layer following live cell time-lapse observation to gain additional biological readouts. We assessed growth arrest induction via cellular status of cell cycle inhibitor p21 four days post irradiation - activated in consequence of individual hit patterns (Figure SI9). First, the correlation of clonogenicity and p21 signal resulted in a binary distribution mirroring the binary readout of p21-positivity and -negativity with assessable threshold intensity (Figure 7c). To further investigate the links of energy deposition, clonogenicity, and induction of growth arrest, we plotted all three parameters for each single cell in one graph and defined quadrants based on p21-positivity threshold and median hit number (Figure 7d). Comparable correlation for dose is shown in Figure SI8. As expected, cells from lineages of high clonogenicity (defined as ≥8 progeny cells originating from a single initial cell) predominantly expressed low (≤ 400 a.u.) p21 nuclear intensity, and rather received initial hit numbers in the lower to intermediate range (10 ≤ hits < 27). Accordingly, respective cell progeny constituted 65 % of the experimental endpoint population. Notably however, above average hit numbers (median = 27) or dose did not necessarily limit clonogenic potential or induce p21, as more than one third of the progeny of high hit cells were determined p21-negative (comparing sector II: 12 % to sector III: 22% of the overall population). This indicates the existence of a subgroup in which carbon ions failed to induce clonal restriction via growth arrest induction. Vice versa, we also determined single cell linages with low clonogenic potential independent of p21, which indicates alternative growth arrest induction mechanisms or asymmetric fate determination of daughter cells. Indeed, we could trace back the lineages of single cells of interest and detect representative cases of differential cell fate determination in terms of subsequent proliferation and p21-expression following cell division (Figure SI2). Taken together, these results demonstrate crucial biological influence on cell fate on an energy deposition-dependent background.

## Discussion

Here, we demonstrated that the design and application of our biomedical sensor *Cell-Fit-HD*^*4D*^ clearly differs from conventional experiments which characterizes cellular response after irradiation on a population level: We are able to determine and to correlate each cells individually received energy deposition by number of hits and dose and to correlate these physical parameters with the temporal and spatial evolution of each cell as well as its molecular kinetics (Figure 6 and 7). We are now able for the first time to access the response variability of a tumor cell line to the inhomogeneous microscopic dose deposition by a clinical ion beam (Figure 4, 5). The implementation of fluorescent reporters to the biomedical sensor allows assessment of various biological parameters in real time such as cell cycle phase or pathway activation as a function of energy deposition. With these findings we were able to start resolving the cell’s individual response chain triggered by the physical energy deposition in its nucleus. This further means that, by forming post-experimental groups of cells with similar deposited energy pattern in *silico*, we can potentially perform experiments within a single run which otherwise require multiple irradiations with altered parameters.

### Design of Cell-Fit-HD^4D^ and its workflow

*Cell-Fit-HD*^*4D*^ was designed to allow for standard cell culture treatment of the biological compartment. This enables a read-out of the biological compartment by standard widefield microscopy, which is identical to the imaging of ordinary multiwell plates. The independent read-out of the cell layer allows for a fast acquisition of multiple imaging fields in live-cell imaging modus with low phototxicity. To record the physical compartment (i.e. FNTD) *Cell-Fit-HD*^*4D*^ is simply mounted on a CLSM. The confocal read-out of the FNTD imaging with relatively high laser power at the end of the live cell imaging does not affect the viability of the cell layer. Despite the decoupling of the read-outs the registration procedure enables the reconstruction of the physical energy deposition of the ions with a spatial accuracy (< 1.5 µm, *paragraph Assessment of error sources in supplementary information*) which is much smaller than the dimension of a single cell nucleus. The independent read-outs of both compartments have the unique potential to expand the field of potential users of *Cell-Fit-HD*^*4D*^. Widefield microscopy is a standard tool in many life-science and clinical research facilities. The read-out of the FNTD can be sourced out to a facility having access to confocal microscopy or the FXR700RG reader (Landauer Inc.) [15]. Since the biomedical sensor can directly be transferred to the microscope after irradiation for read-out the risk of contamination of the cell layer (including the culture medium) is minimized.

The design of *Cell-Fit-HD*^*4D*^ shows promise for cell coating with 3D cell culture or even human biopsies in order to reflect better the *in-vivo* situation of tumor treatment by ion beam therapy. The biological compartment can principally be imaged without any fluorescent staining (brightfield) simplifying the modification and workflow of the biopsy. In addition, the simple and cost effective design of our biomedical sensor allows its usage by laboratories with limited access and expertise in detector technologies or microbeam delivery systems.

To overcome the limitations (given by the excitation and emission spectra of the FNTD [10]) in using fluorescent dyes in the cell layer with a wavelength spectrum in the blue and far-red, region we developed an alternative Mounting-architecture (Figure SI3a). There, the FNTD is not in direct contact with the monitored cell layer. A sufficient distance between both compartments was introduced to avoid interference of the read-outs of the physical- and biological compartments and to allow the usage of the full wavelength spectrum of fluorescent dyes. The Mounting-architecture is described in detail in the *Supplementary Information* (*paragraph Mounting-architecture architecture*).

A short time interval between irradiation and initial read-out (step 1 in Figure 3a) is beneficial for a precise computation of the energy deposition by ions in individual cell nuclei. A large time frame could be accompanied by significant cell migration and hence false assignment of ion traversals to individual cell nucleus. Despite a short time interval between irradiation and imaging, of approximately 5 minutes, we did not attempt to correlate individual ion traversal with its nearest neighbour 53BP1 focus. We expect a significant risk of false correlation using the irradiation dose of 1 Gy with its median number of 27 ion traversals in a single cell nucleus (Figure 5). In *Supplementary Information* (paragraph *Assessment of error sources/ uncertainties in spatial correlation*) we listed all parameters in the workflow of *Cell-Fit-HD*^*4D*^ which could affect the spatial correlation of physical energy deposition and cellular response. Additionally, we introduced a Monte Carlo approach to principally simulate the error propagation. Using this approach we computed the the mean total error to be much smaller than the dimensions of a cell nuclei (cylinder of diameter 10 µm).

The interval *Δt* between two consecutive time points in the live-cell imaging (step 2, Figure 3a) can principally be varied. We applied the combination of *Δt= 15 min* (first 9 time points) and *Δt= 45 min* to record the early DNA damage repair but also to avoid phototoxicity by the widefield imaging at later time points. A sole recording of the cell layer by bright field – without fluorescence imaging – could further decrease the photo toxicity and hence shorten *Δt*.

The cell seeding density and hence the number of cells to be imaged can principally be adapted according to the aim of the experiment. The initial stack (step 1 in Figure 3a) is the major factor influencing the initial number of cells to be imaged. The recording of the initial stack for a single imaging field lasts approximately 30s. This acquisition time could principally be reduced by a declined number of imaging planes in the initial stack. Increasing the number of initial stacks would also increase the time interval between irradiation and initial recording of the cell response to irradiation. As mentioned above, this can lead to increased miss-assignments of ion traversals to the corresponding cell nuclei. Furthermore, the initial stack is crucial to account for the uncertainty in stage movement of the widefield microscope (*paragraph Precision of microscopy stage movement, Supplementary Information*). However, the new design and corresponding workflow of *Cell-Fit-HD*^*4D*^ drastically increased the number of cells analysed after irradiation comparing to former designs of *Cell-Fit-HD* in which only confocal read-out was possible [24].

### Correlation of individual molecular and cellular dynamics and deposited energy

Measurement of radiation-induced DNA-damage foci by immunostaining of respective proteins presents a widely accepted standard for dose assessment. Such analysis is mostly performed on fixated cells and dose correlations are conducted with average population numbers. However, as in ion beam therapy a homogenous irradiation field is not applied on the microscale level (Figures 4, 5), this method cannot describe a true dose-effect relationship. The biomedical sensor presented here allows assessment of the relationship of deposited energy and resulting DNA damage foci in single living cells over time. Our analysis on single cell level revealed a principal linear correlation between energy deposition and RIF-occurrence which was obscured by broad biological and individual cellular response (Figure 6, SI6).

To better understand RIF formation dynamics in ion fields, we performed split analysis for received hits (irrespective of LET) and dose (irrespective of hit number). Here we show that for a cell mono layer, the LET of single ion traversals in the cell nucleus has lesser impact on RIF number than the total number of intranuclear ion hits, as implicated by superior correlation. At equal dose, high numbers of lower LET ions create higher RIF numbers compared to low numbers of high LET ions (Figure 8). This however might change for wider LET spectra and RIF integration for a single ion track passing cells in a multilayer.

**Figure 8.**
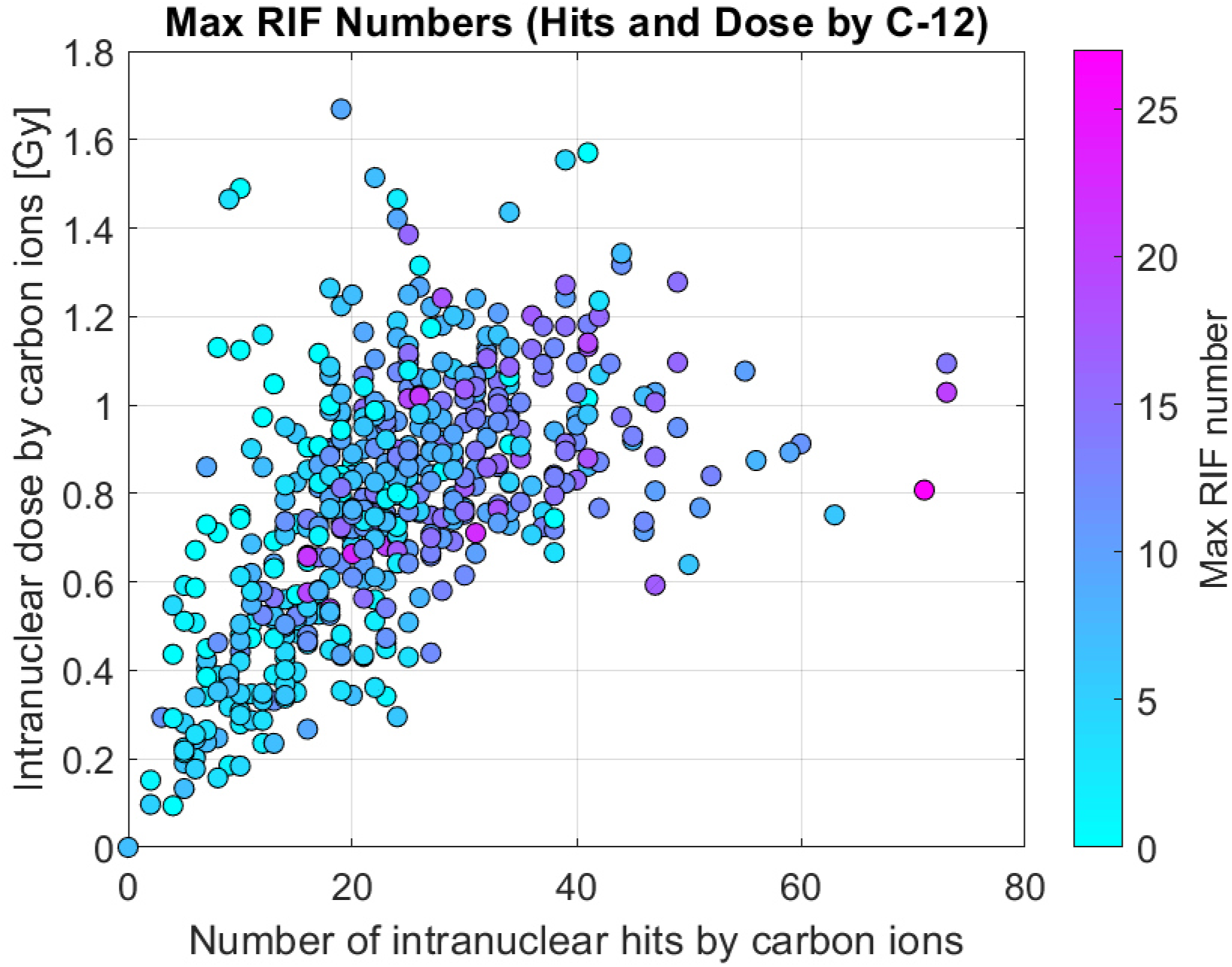
The number of ion traversals in a nucleus has an impact on formation of RIFs. At equal dose, a high number of hits by primary carbon ions (C-12) of low LET seem to create more RIFs (maximum RIF numbers within 25 h post irradiation) than a low number of hits of high LET. Planned physical dose of 0.5 and 1 Gy. N= 514.

While prominent time points for RIF assessment usually are 15 min, 1h, or 24h post irradiation, we found maximum linearity in the interval between 5.6 h and 10.9 h post-irradiation (Figure 6b). We suggest this might reflect a kind of balance state in which the ratio of established RIFs and already repaired RIFs reaches a maximum. Ongoing occurrence of RIF at this time point is indicated by an increasing progression slope, while completed repair becomes apparent by general reduction of RIF numbers after ∼ 50 min (Figure SI1). In conclusion, static RIF assessment at time points with peaking RIF numbers do not necessarily mirror received energy in the best way.

In contrast, we could show that dynamic RIF monitoring and ascertainment of maximum RIF numbers in a cell nucleus within 25 h after irradiation increases the power as a surrogate and potential biomarker for deposited energy by ions in individual tumor cell (Figure 6d, 9) [28]. The correlation of maximum RIF numbers with hit numbers increases the Pearson-r coefficient to 0.47. Furthermore, we proposed the physical parameter ΣLET (sum of the ions’ LET in a cell nucleus) to provide an additional quantification /alternative estimation of the ion’s impact on the biological response. The application of this parameter for correlation with maximum RIF numbers further elevated the Pearson-r coefficient as compared to RIF/dose correlation, and is in the range of the coefficient for hit number. This points towards a certain independency of the molecular and biological impact of single ions from the target volume in this 2D cell layer setup, in which the single cell nucleus presents the most basic biological entity.

**Figure 9.**
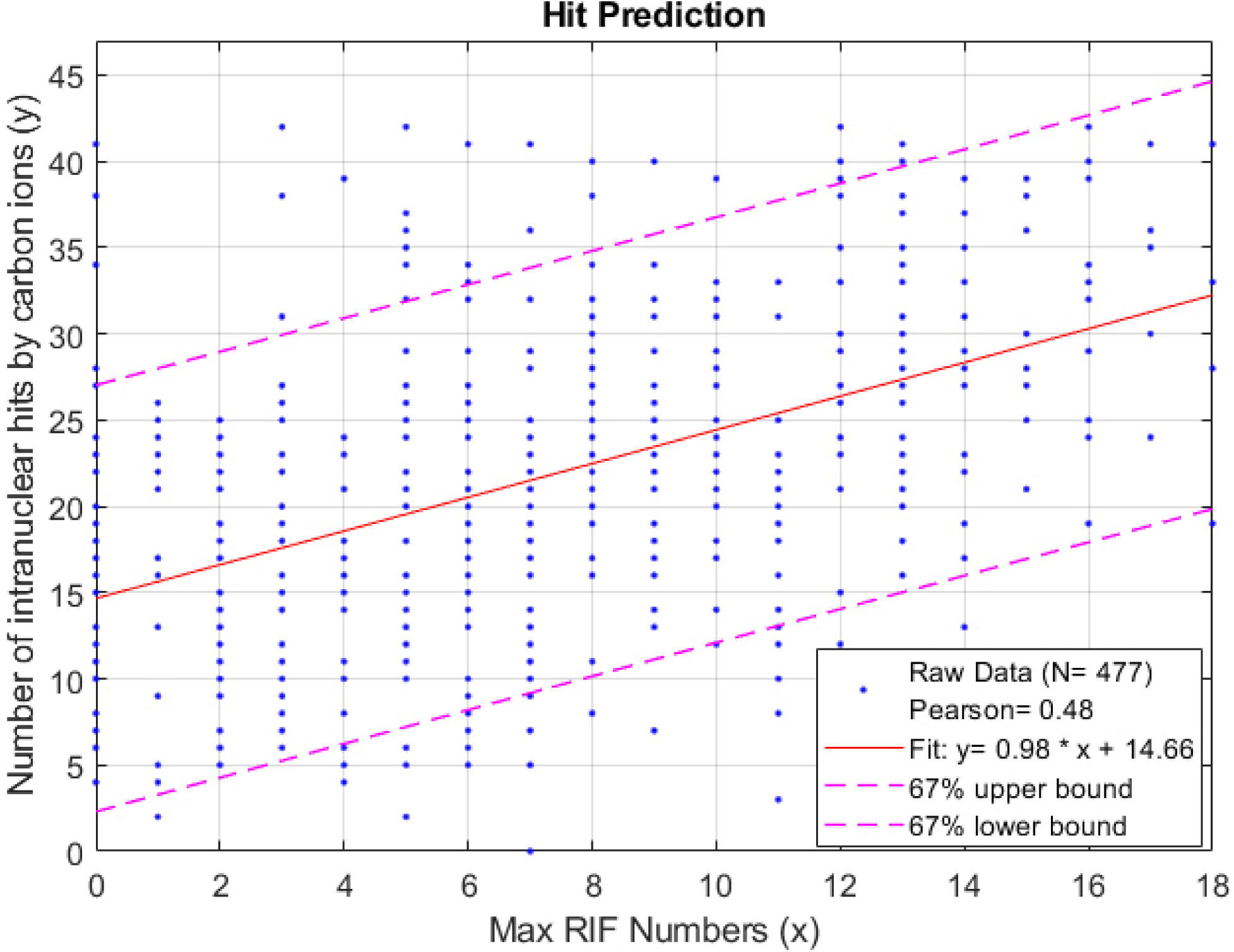
Maximum number of RIF (53BP1 foci) within 25 h after irradiation is a potential biomarker for the physical energy deposition in the cell nucleus. There is a linear correlation (red line) between the maximum RIF numbers and the number of ion hits by the primary ions (carbon ions, SOBP, planned physical dose of 0.5 and 1 Gy). Confidence intervals (67%, magenta dashed lines) are depicted. The upper bound of the maximum RIF number of 18 was chosen to maximize the Pearson-r-coefficients.

By the determination of maximum RIF number at a certain time point, only RIFs simultaneously existent within a cell are measured and, consequently, already resolved RIFs will not be included. Potential RIF tracking or total summation of all emerging RIF within a cell is not likely to improve RIF detection due to over-interpretation of signalling noise and the fact that 53BP1-foci vanish prior to mitosis [29, 30]. Quantification improvement however should be feasible with stabilization of construct expression and switch to reporter proteins remaining on DNA damage during mitosis like MDC1[29].

As mentioned earlier, the described phenomena in RIF formation were observed on a background of immense heterogeneity and variable cellular responses. This implies a vast dependence of RIF dynamics on physical, molecular and biological pre-requisites. First, the ions’ ionization pattern originates from stochastically occurring events and determines initial damage infliction and its complexity. Next, RIF formation occurs in a delayed fashion in regions of densely packed heterochromatin, and especially in cancer cells, the general grade of chromatin packaging may vary substantially between subclones [31, 32]. In a similar fashion, repair capacity is highly dependent on DSB accessibility for respective proteins, an aspect which is linked to chromatin state as well as damage complexity [33, 34]. In general, DSB repair is facilitated by a highly complex molecular machinery which is regulated on numerous levels. Therefore, repair kinetics are influenced by cell-specific levels of factors (proteins, micro-RNAs) which regulate repair pathway activity at the time point of irradiation [35-37]. It is further known that dynamics of DNA damage foci emergence and repair are altered with ongoing cell cycle phase due to specific regulation of repair pathways like non-homologous end-joining (G1) and homologous repair (G2) [38, 39]. Such aspects help to explain the ion hit/dos-independent variability in RIF formation observed in this study (error bars in Figure 6b and SI6b). The future combination of *Cell-Fit-HD*^*4D*^ with additional fluorescent tags and reporters will further help decipher RIF formation in response to cell-specific energy deposition.

While others have shown multiple RIFs formed along single ion traversal through nuclei within sub-micron distances [40] we did not detect a RIF for every scored ion track traversal in the z-projected images (∼ 0.2/0.23, Figure 6b,d). It has to be emphasized that the RIF induction for individual hit numbers is subjected to great fluctuations (Figure 6b). For certain hit numbers the actual RIF/Hit ratio can be much higher, e.g. 0.9 (Figure SI6e). The probabilistic nature of ionization events can lead to ions passing the nucleus without damaging the DNA significantly enough to trigger the formation of any RIF along the ion trajectory. This has been predicted and observed for different ion species and LET values [26]. Heiß et al. [41] irradiated cells with pre-defined geometrical pattern of single ions, but only observed a clearly corresponding RIF pattern in 61% of the cases. In their setup, carbon ions with energies of 4.8 Mev/u (∼290 keV/µm), which represent multi-folds of the LETs applied in our experiments, were applied. The fact that RIF formation is principally dependent on LET further helps explaining the apparently low ratio of RIF/hit with the clinically-relevant energies administered here. We exclude substantial false-negative detection of RIF based on the results of our validation trials as well as the large, signal intense nature of heavy ion induced RIF. Nonetheless, widefield microscopy with a 20x objective has restrictions regarding resolution, so that we cannot fully exclude additional presence of short-lived, low intensity RIF. However, these are usually attributed to have lesser biological relevance and are even more neglected in standard, static RIF measurements in fixed cells due to lacking temporal resolution.

Regarding RIF induction dependency on the single cell level, a higher correlation was found with ion hit numbers compared to deposited dose (Figure 6a). This reflects that microscale dose is the sum of distinct LETs of single ions traversing the cell. Therefore microscale dose can attain equal values by various combinations of hit numbers and LETs. For RIF-hit relationships, one of these parameters is fixed, what probably leads to lesser variability for distinct hit numbers of cells. In other words, while variability in RIF-hit relationships is influenced by one physical parameter, namely LET-distribution, RIF-dose relationship is affected by two, that being ion hits and their respective LET.

The data therefore leaves room for speculation that, within the here applied LET range, the probability for induction of a DNA damage entity by an additional ion traversal is higher as for LET increase of an existing one. This however might be valid only without regard to the quality of damage, as we did not regard RIF cluster sizes on the z-level. Generally, *Cell-Fit-HD*^*4D*^ presents a suitable tool to further shed light on this topic [24, 42].

The measured number of RIFs per ion traversal (∼ 0.2/0.23, Figure 6b, d) in this work is principally well in line with *in silico* simulation of RIF induction by Monte Carlo approach using Geant4-DNA. There, simulated RIF formation was compared to 53BP1 foci detected per proton or alpha ion track [27]. Despite this good agreement the hypothetical question remains why not at least one 53BP1 focus along each ion traversal in a cell nucleus (thickness ∼ 4 µm) is detected in our presented work. It is important to underline that the RIF induction for individual ion was not determined here despite *Cell-Fit-HD*^*4D*^ being an appropriate tool for this purpose. The RIF induction along single ion traversal is principally LET-dependent [40]. A movement of cell nuclei within the short time interval (∼ 5min) between irradiation and imaging (Figure 3) and the clinical-relevant ion beam fluence (median hit number ∼ 27 primary hits, 1 Gy SOBP) could alter the hit-RIF-assignment. Instead, the total number of intranuclear ion hits of wide LET distribution (Figure 4b) was used to calculate a mean RIF induction ratio for a cell nucleus. The LET-dependence of RIF induction is basically reflected in the error bars (SD) in the hit-RIF-correlation (Figure 6b). Each error bar represents a LET distribution for each hit number. The broad LET distribution with increasing hit numbers contributing to a given dose could also be the reason for the stagnation of the slope in linear regression in the RIF-dose correlation (Figure SI6b). A possible reason for not detecting at least one RIF per ion hit could be short-living RIFs of low 53BP1 signal, which could not be detected by the *Trainable Weka segmentation* plugin. In our verification measurements (*paragraph γ-H2AX vs endogenous 53BP1, Supplementary Information*) we however determined a spatial overlap of ∼ 74% of γ-H2AX foci (gold standard in DSB detection) and 53BP1 foci as well as the detection of large 53BP1 foci originating from complex DSB clusters being induced by carbon ions. Generally, *Cell-Fit-HD*^*4D*^ and microbeam technology are future tools to further shed light on this topic [24, 42-44].

In order to assess biological effectiveness of irradiation, primary cell lines like Chinese Hamster Ovary cells or fibroblasts are often taken as reference. For cancer cells, however, irradiation outcome on the cellular level may potentially vary substantially at fixed microscopic energy depositions since they circumvent and downregulate cellular shutdown programs like apoptosis and senescence [45, 46]. The presented biomedical sensor enables for the first time decryption of true dose/hit-response relationships up to several days (Figure 10). Within this period of observation, restricted numbers of cell division events can be observed. By low cell number seeding it should be feasible to determine to what extent clonogenic potential within the first couple of days upon irradiation correlates with formation of “classical” colonies, i.e. 50 cells of progeny. The data obtained in this work can be interpreted as a typical scenario in which proliferative potential of cancer cells is restricted in a dose/hit-dependent manner, with single cells escaping growth arrest despite receiving high number of hits (Figure 7d). Notably, *Cell-Fit-HD*^*4D*^ allows further characterization of such cells in terms of morphology or active cellular programs via fluorescent reporters or endpoint staining, as we demonstrated for p21 (Figures SI2, SI9). It is also possible to trace back their lineage to test for potential asymmetric cell division with daughter cells undergoing differential fate, as we observed frequently (Figure 7a). Therefore, *Cell-Fit-HD*^*4D*^ presents a powerful tool to fully comprehend the mechanisms of ion radiation-resistance in an energy-pattern-resolved manner.

**Figure 10.**
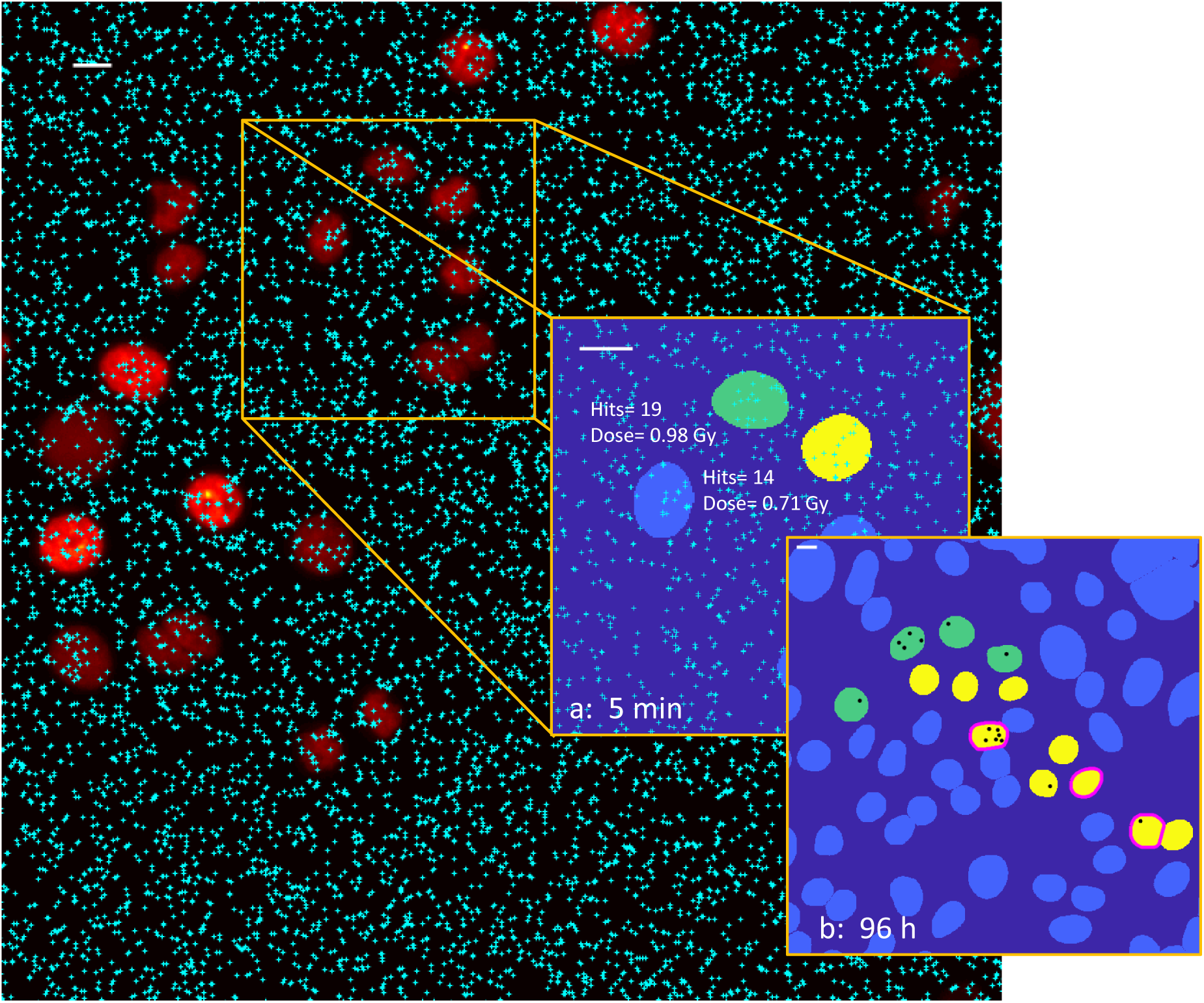
*Cell-Fit-HD*^*4D*^ combines single-cell dosimetry with molecular and cellular dynamics of individual tumor cells. Here we exemplarily demonstrate the correlation of number of intranuclear carbon ion hits (irradiation with planned physical dose of 1 Gy) with the time and space evolution of each cell and its molecular kinetics. Fluorescent signal of 53BP1 damage protein in A549 cell nucleus is shown. Each cyan cross corresponds to a single reconstructed ion traversal. The cross-sectional areas of segmented cell nuclei are depicted in (**a**) and (**b**). A set of mother cells in (**a**) were monitored and tracked within 96 h after irradiation. Daughter cells in (**b**) are shown with the same color as their mother cell in (**a**). **(b)** Persistent 53BP1 foci (RIF, black objects) and p21 status of each cell 96 h after irradiation are displayed. P21 positive cells (p21 signal ≥ 400 a.u.) are encircled in magenta. Scale bars, 10 µm.

## Conclusion and Outlook

Here we present the biomedical sensor *cell-fluorescent ion track hybrid detector*^*4D*^ (*Cell-Fit-HD*^*4D*^) which enables long-term monitoring of single tumor cells after clinical ion beam irradiation in combination with single-cell dosimetry. In contrast to existing radiation experiments, our sensor combines individual monitoring and tracking of a large number of cells over several days by conventional widefield microscopy and visualization of the physical energy deposition in cellular compartments (Figure 10). This sensor enables for the first time combined visualization of a clinical carbon ion field and the spatiotemporal correlation of single ion traversals with the response on subcellular and cellular level.

These unique properties of *Cell-Fit-HD*^*4D*^ allow translation of the heterogeneous microscopic energy deposition of ions into the individual (and variable) cellular response (Figure 11). The three-dimensional correlation of intranuclear ion traversals, p21 expression and clonogenicity is just one example of such translation (Figure 7d). It helps to address the following question: Which combination of biological cell parameters (e.g. repair dynamics, proliferation) and physical irradiation parameters (e.g. number of cell hits, dose, dose rate) does a radiation-resistant tumour cell exhibit? In addition, our analysis (Figure 6) identified the number of intranuclear ion traversals to be a good predictor for the 53BP1 RIF formation (and better than the intranuclear dose, Figure SI6). Therefore, the number of hits could be additional parameter - or even in combination with dose and the proposed parameter ΣLET - to quantify the cellular response on molecular and population level. Our biomedical sensor could provide crucial input for current mechanistic approaches to biophysical modelling of the effect of ionizing radiation to biological matter.

**Figure 11.**
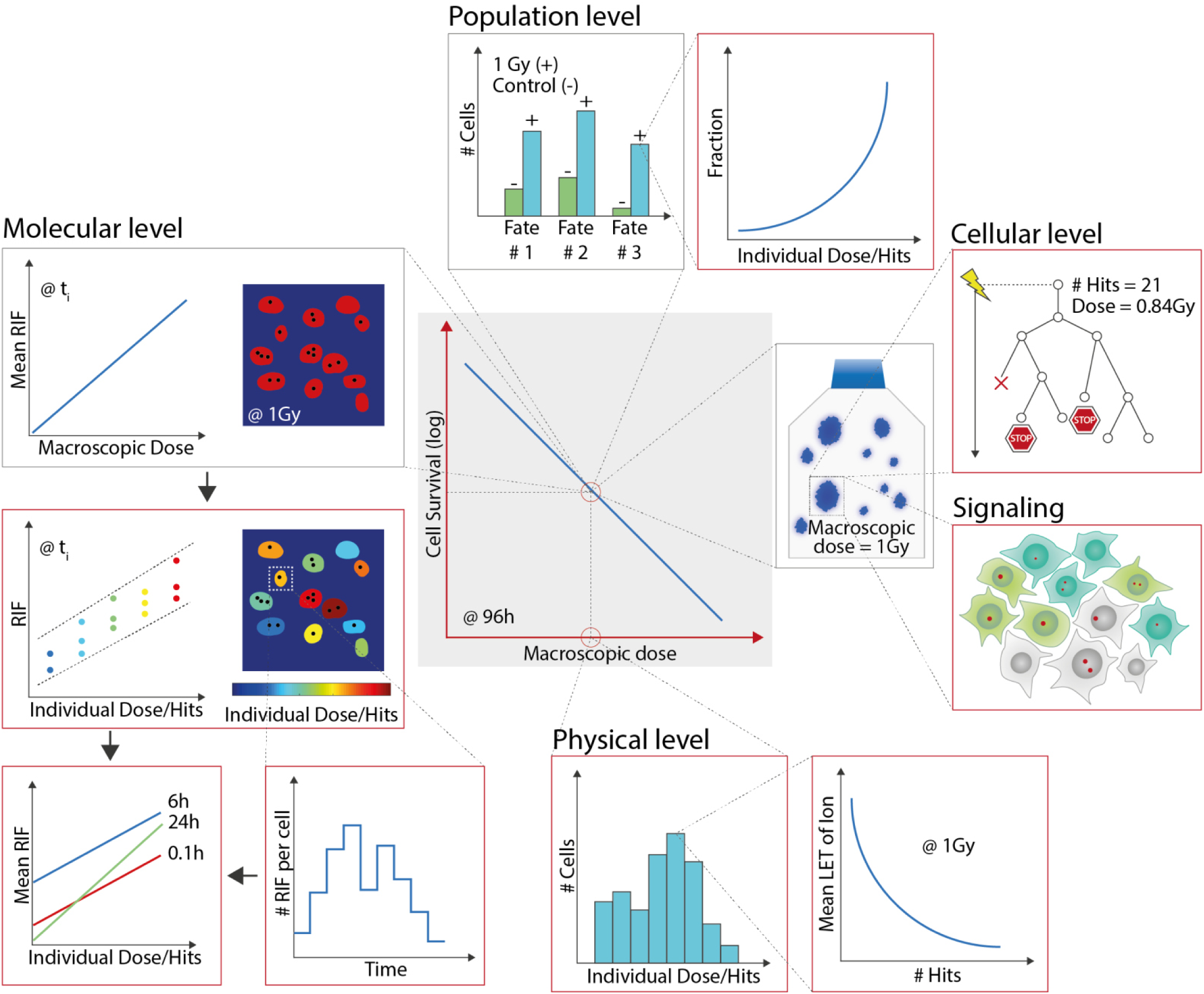
Schematic illustration of current standard radiobiological experiment (framed in black) and beyond deciphered by *Cell-Fit-HD*^*4D*^ (framed in red). Cell-survival-curve (blue solid line, grey box) presents the current gold standard in defining the effect on irradiation outcome and effect. In standard cell culture irradiation it is assumed that each cell is facing identical dose in a macroscopic ion field, termed macroscopic dose. By *Cell-Fit-HD*^*4D*^ the physical energy deposition for individual cell nucleus is however resolved, termed individual dose/hits. LET: linear energy transfer. **Molecular level:** In contrast to a mean RIF value at time point *t*_*i*_ for a certain macroscopic dose value, *Cell-Fit-HD*^*4D*^ enables to resolve the time-dependent RIF evolution as a function of individual dose/ hit deposition in a cell nucleus (colour coding). For each dose/hit value occurring in a microscopic ion field the RIF distribution is assessed. **Cellular/Population level:** The binary readout of existing or non-existing colonies is extended by clonal dynamics and clonal fate, providing biological information on how the colony emerged. In common cell survival experiments different cell fates are assigned to the macroscopic dose value. Using *Cell-Fit-HD*^*4D*^ sub-fractions of a cell fate is resolved according to the individual dose/ hit deposition in the cell nucleus. **Signaling:** Additionally, the landscape in terms of signalling between individual daughter cells derived from a mother cell can be assessed by *Cell-Fit-HD*^*4D*^.

In the clinical context, providing multi-dimensional physical and biological information on individual tumor cell is an important step for determination what tumor entities can be effectively treated by ion beam therapy and for understanding local recurrence of a tumor after ion irradiation. The present treatment planning underlying physical beam parameters could be extended by a set of novel physical (e.g. number of intranuclear hits) and biological (e.g. cell cycle and signalling or immune response) parameters to maximize tumor control and minimizing normal tissue complication. New target points in the cellular response cascade to ion irradiation can be detected in order to create multimodal – i.e. combination of ion beam therapy with drug application - oncological concepts to maximize the benefit for the patient.

*Cell-Fit-HD*^*4D*^ is an important tool for the search of new biomarkers for indirect visualisation of the physical dose deposition (Figure 9). Design of *Cell-Fit-HD*^*4D*^ gives promise for cell coating with 3D cell culture or even human biopsies in order to better investigate the *in-vivo* (real) response to tumor irradiation.

## Acknowledgements

The authors thank K. Rein for her technical expertise and support in creating illustration. The authors thank A. Mairani and S. Mein for the Monte Carlo (FLUKA) simulation of the ion irradiation and support for the irradiation planning at the Ion-Beam Therapy Center (HIT) of Heidelberg University Hospital; S. Brons for generously providing support and technical irradiation assistance at HIT. The authors also thank F. Bestvater and D. Krunic of the light microscopy core facility at the German Cancer Research Center for their enthusiasm and unflagging support.

## Supplementary Figures

**Figure SI1.**
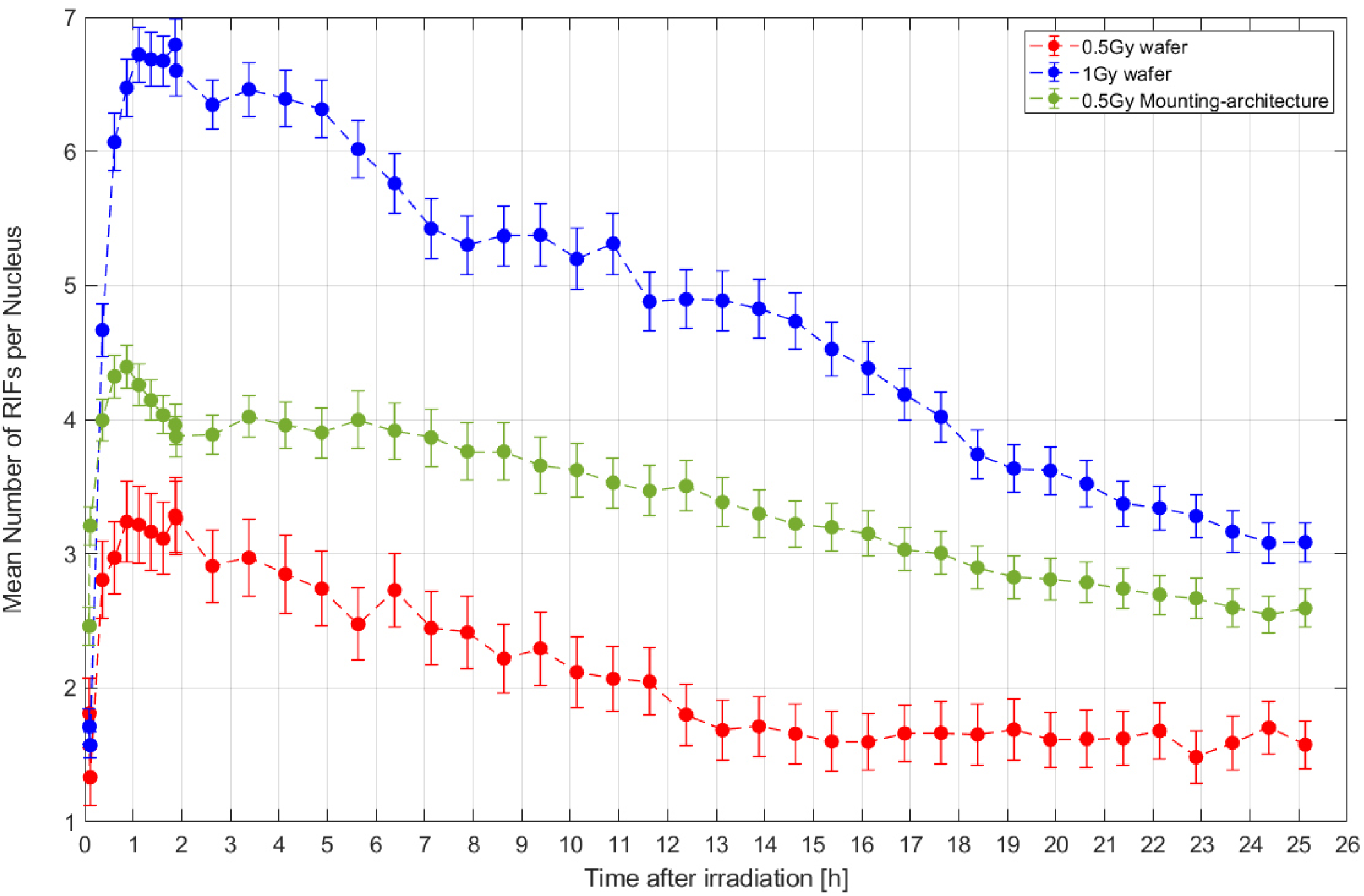
Time dependent behaviour of the mean number of RIFs in a nucleus. Cells exhibit maximum RIF numbers approximately one h after irradiation. The RIF dynamics is shown for irradiated cells cultured on the FNTD surface (red: 0.5 Gy, C-12, SOBP, blue: 1 Gy C-12, SOBP) and using the Mounting-architecture (green, 0.5 Gy, C-12, SOBP). Error bars are the SEM. Time point zero corresponds to irradiation.

**Figure SI2.**
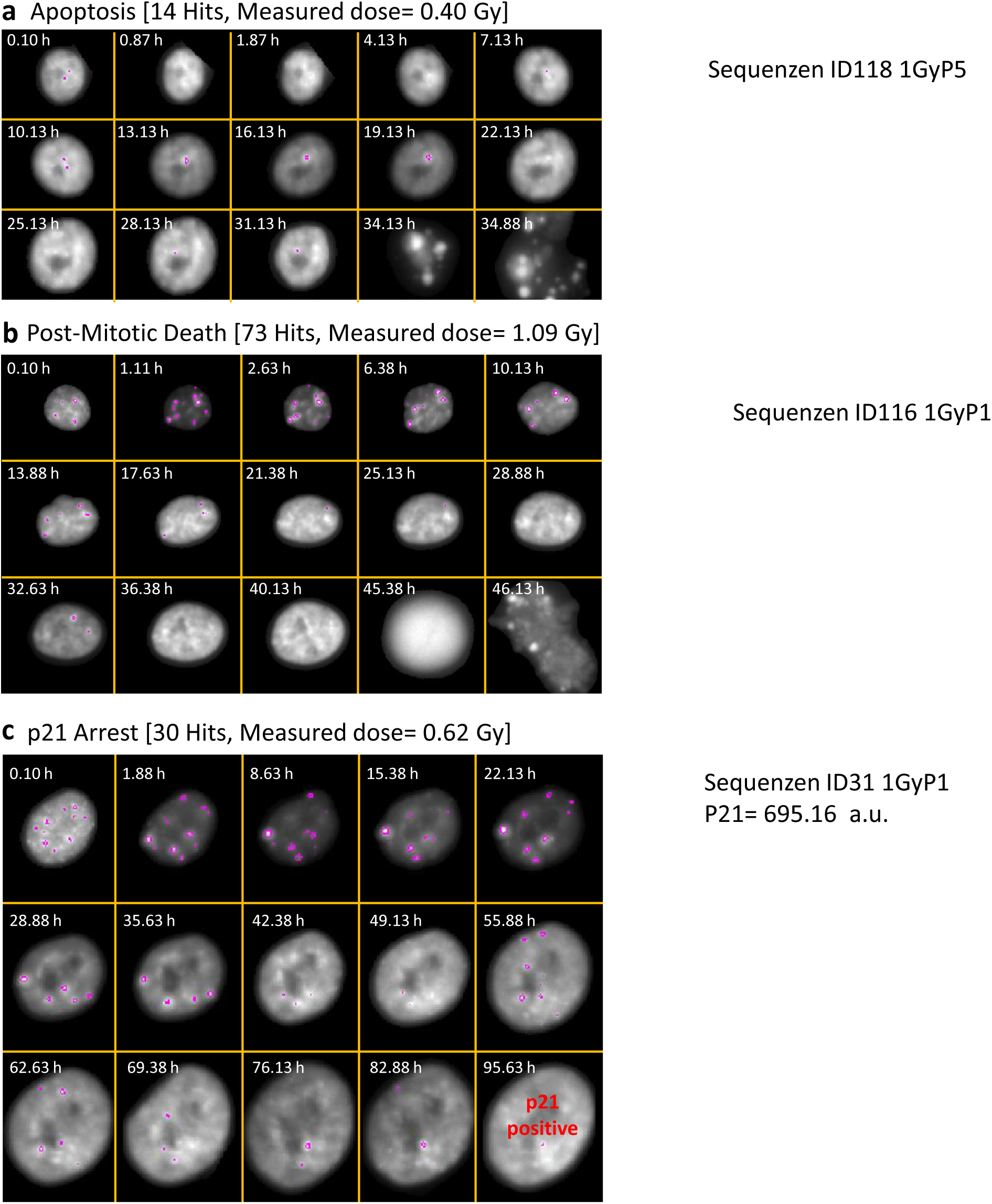
*Cell-Fit-HD* enables online monitoring and tracking of individual cells over several days after ion irradiation. **(a-c)** Visualization of the cellular and molecular kinetics of three exemplary cells after carbon ion irradiation with planned physical dose of 1 Gy. The fluorescent signal of 53BP1 in A549 cell nucleus was recorded over four days by widefield microscopy and is shown here (maximum intensity projection). The RIFs (53BP1 foci) emerging were detected by *trainable Weka segmentation* plugin for *ImageJ* and are encircled in magenta. The tracking and segmentation of individual cells was conducted using *LSetTracker*. The status of the cell-cycle inhibitor p21 was determined by fluorescent staining 96 h after irradiation. For each cell the set of actual physical beam parameters, i.e. physical dose and number of intranuclear hits by carbon ions, is stated in brackets.

**Figure SI3.**
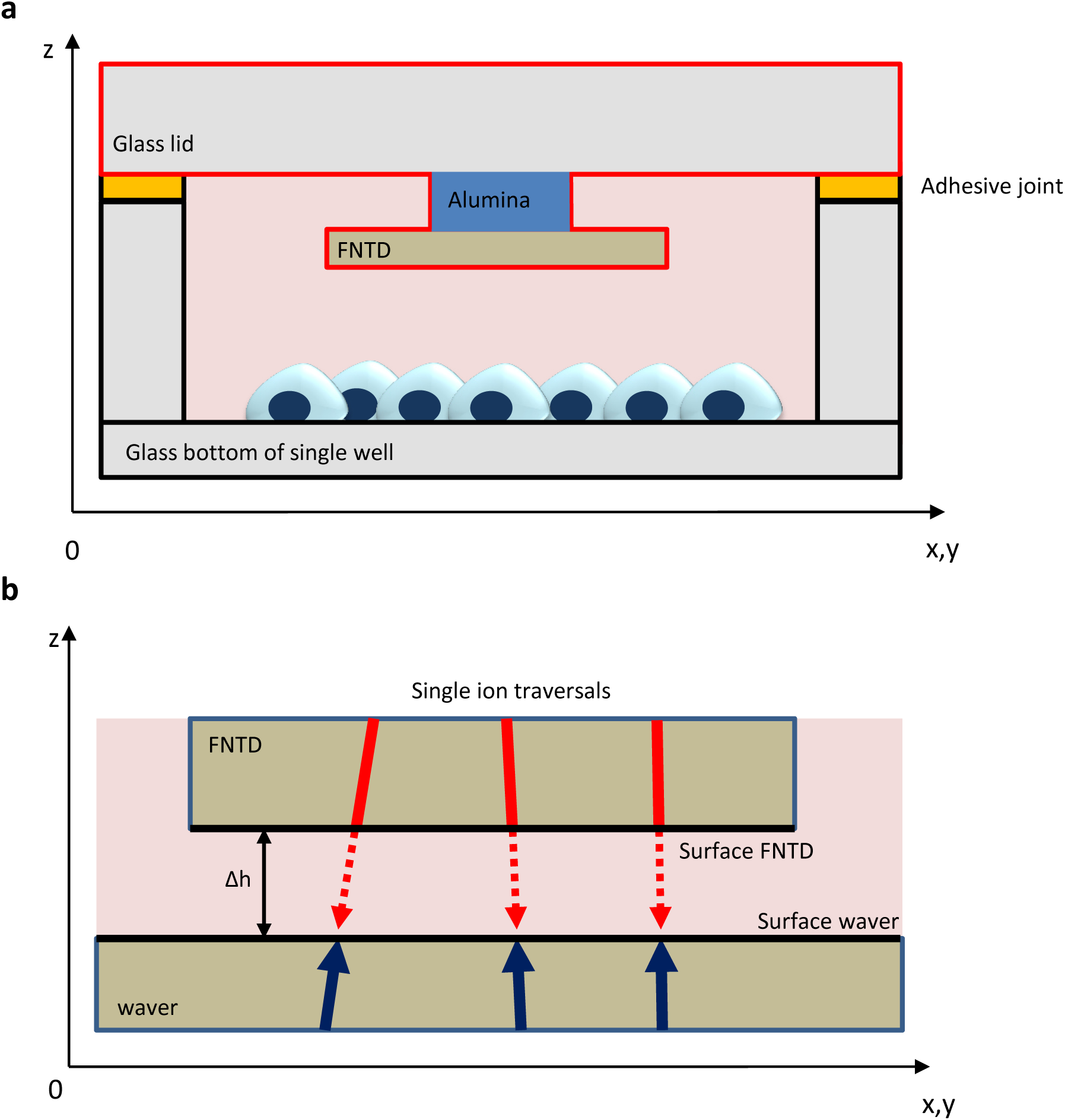
Mounting-architecture, an alternative design of *Cel-Fit-HD*^*4D*^. **(a)** A well of the 12-well-glass bottom dish was covered with a lid containing the FNTD. After the initial read-out, the lid is removed by gently applying force at the adhesive joints. **(b)** Validation of the Mounting-architecture: The bottom of the Mounting-architecture was replaced by a FNTD in form of a thin wafer. After carbon-ion irradiation ion traversals were reconstructed in the FNTD and extrapolated (distance Δh) onto the surface of the wafer. Additionally, identical ion tracks were reconstructed in the wafer. The spatial overlap to the tracks originating from the wafer and from the FNTD were tested.

**Figure SI4.**
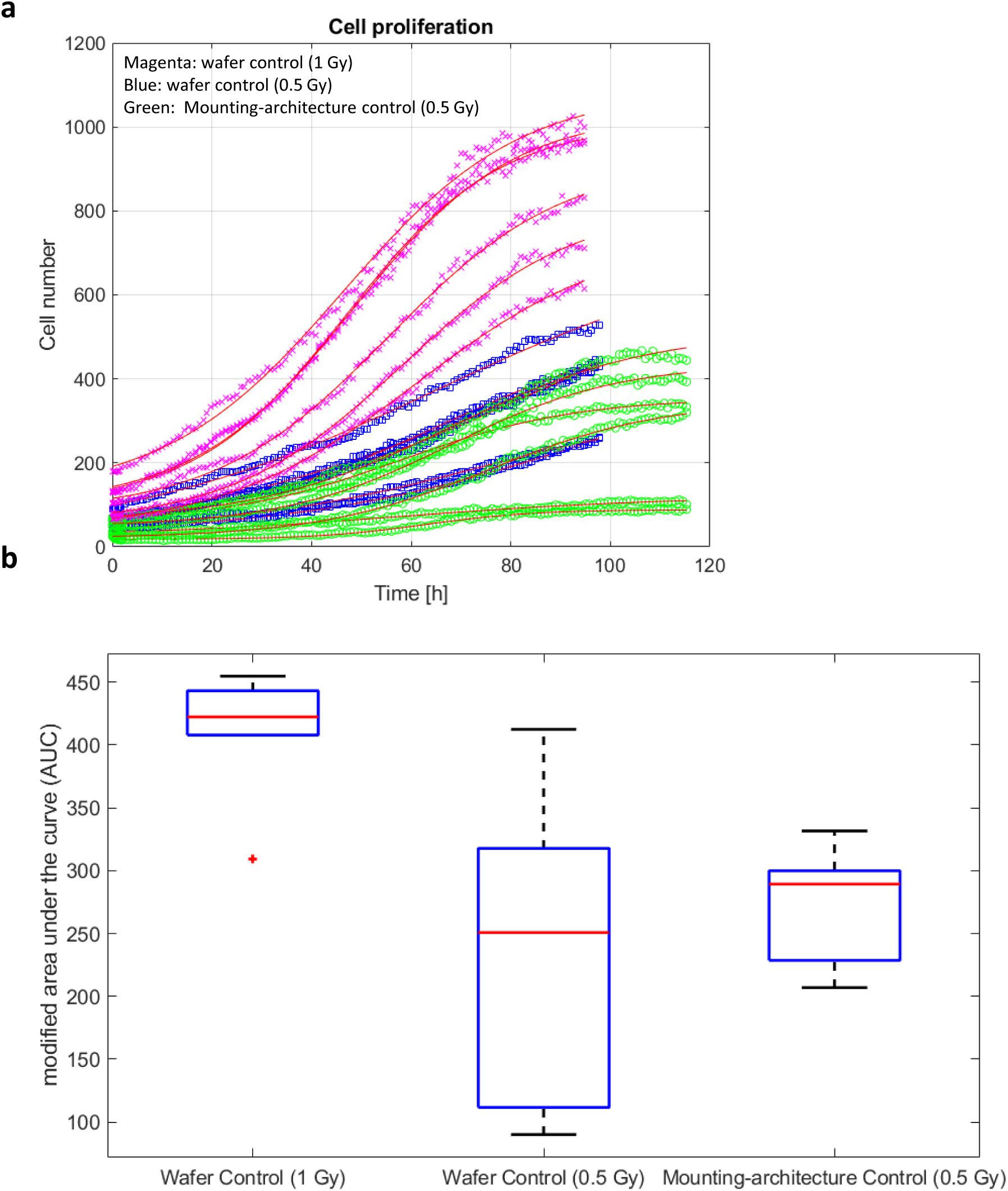
Proliferation curves of cells cultured on wafer and on glass bottom. **(a)** Cell number dynamics was recorded for the controls of the irradiations (magenta: wafer 1 Gy, blue: wafer 0.5 Gy, green: Mounting architecture 0.5 Gy) within a time interval of several days. Six imaging fields were recorded for each control group. Cell dynamics was fitted by logistic function (solid lines). **(b)** Computation of the area under the curve (AUC) in (**a**). For normalization, the number of cells in the initial time frame (time point zero) was used. Medians (red lines), interquartile range (bottom and top blue edges), minimum maximum (black lines), outliers (red cross).

**Figure SI5.**
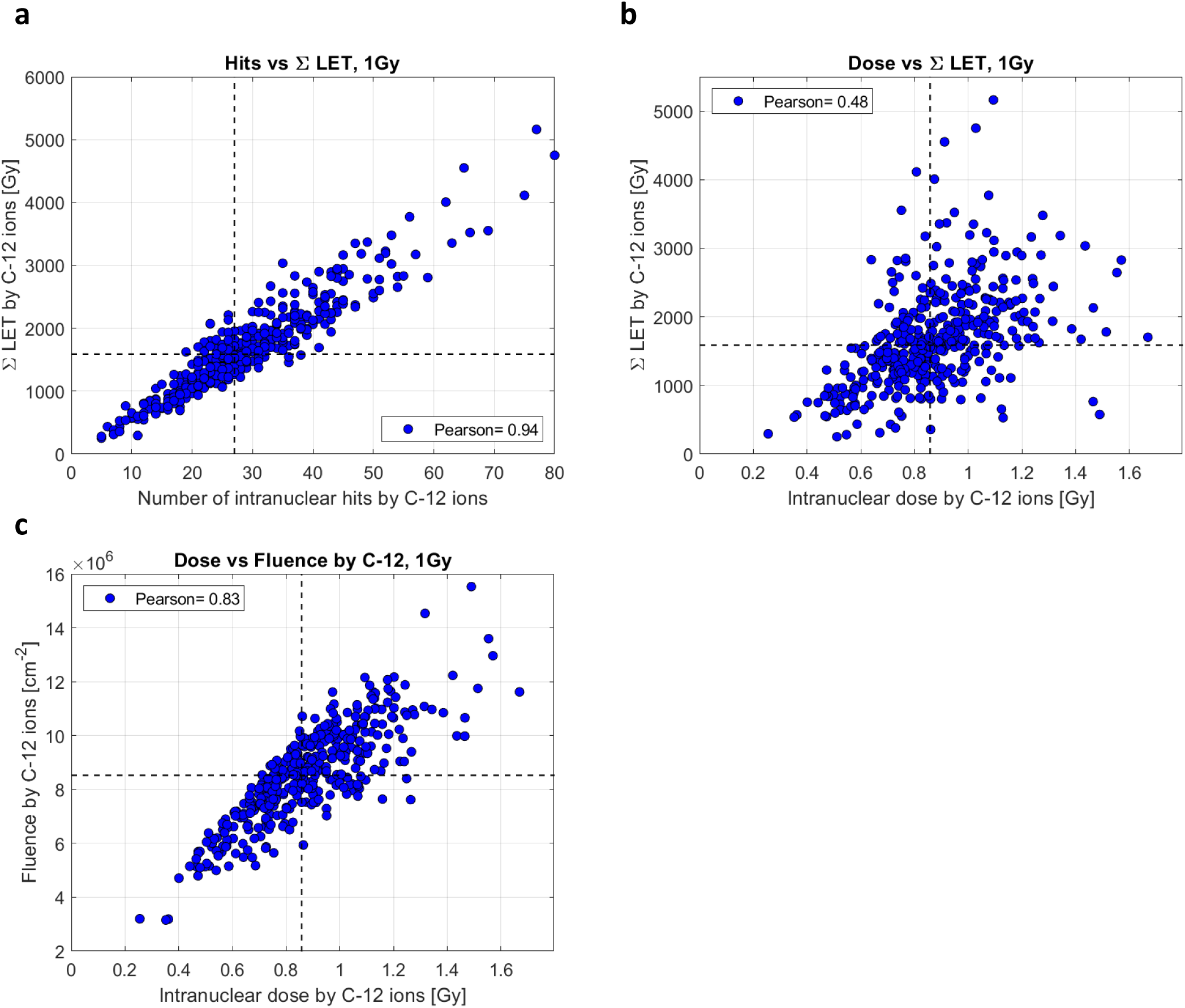
*Cell-Fit-HD*^*4D*^ enables to resolve the distribution of the physical beam parameters on the cellular level. Additional physical beam parameter ΣLET (sum of the ions’ LET in the cell nucleus) was introduced. This parameter disregards the nucleus area and therefore accounts for the single cell as the most basic biological, integer unit. **(a), (b)** Correlation of number of intranuclear ion hits and dose by primary carbon ions (C-12, 1Gy, SOBP) and ΣLET. **(c)** Correlation of dose and fluence by primary C-12 ions. The Pearson-r-coefficients is stated for each correlation. The dashed lines indicate the corresponding median values and were used to define the quadrants.

**Figure SI6.**
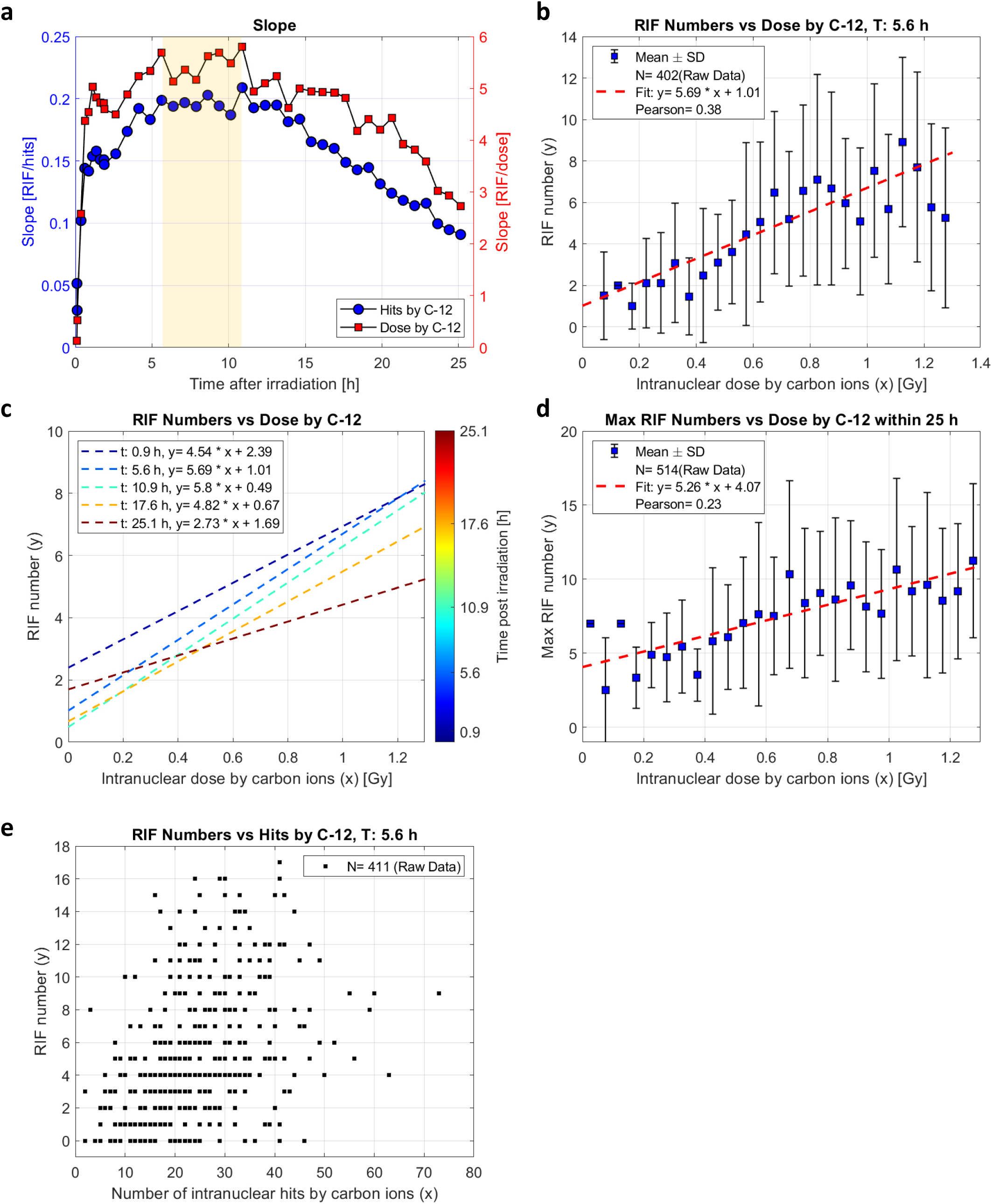
*Cell-Fit-HD*^*4D*^ enables correlation of physical ion beam parameters with molecular dynamics of individual tumor cells. **(a)** Time-dependent slope of the linear regression within 25 h post irradiation. The maximum is in the interval [5.6; 10.9] h post irradiation (yellow bar). **(b)** Linear correlation between the intranuclear dose by carbon ions (C-12) and the number of induced RIFs 5.6 h after irradiation. **(c)** Time-dependence (0.9, 5.6, 10.9, 17.6, 25 h post irradiation) of linear correlation between dose by C-12 and the number of induced RIFs. **(d)** Correlation of maximum number of emerging RIF within 25 h after irradiation with dose by C-12. **(e)** Raw data for the correlation of hits with RIF. For the computation of the Pearson-r correlation and linear regression analysis the raw data was used. Here, the dose was limited to 1.3 Gy in (**a-d**) to ensure sufficient number of data points. Only mother cells were considered in the data analysis.

**Figure SI7.**
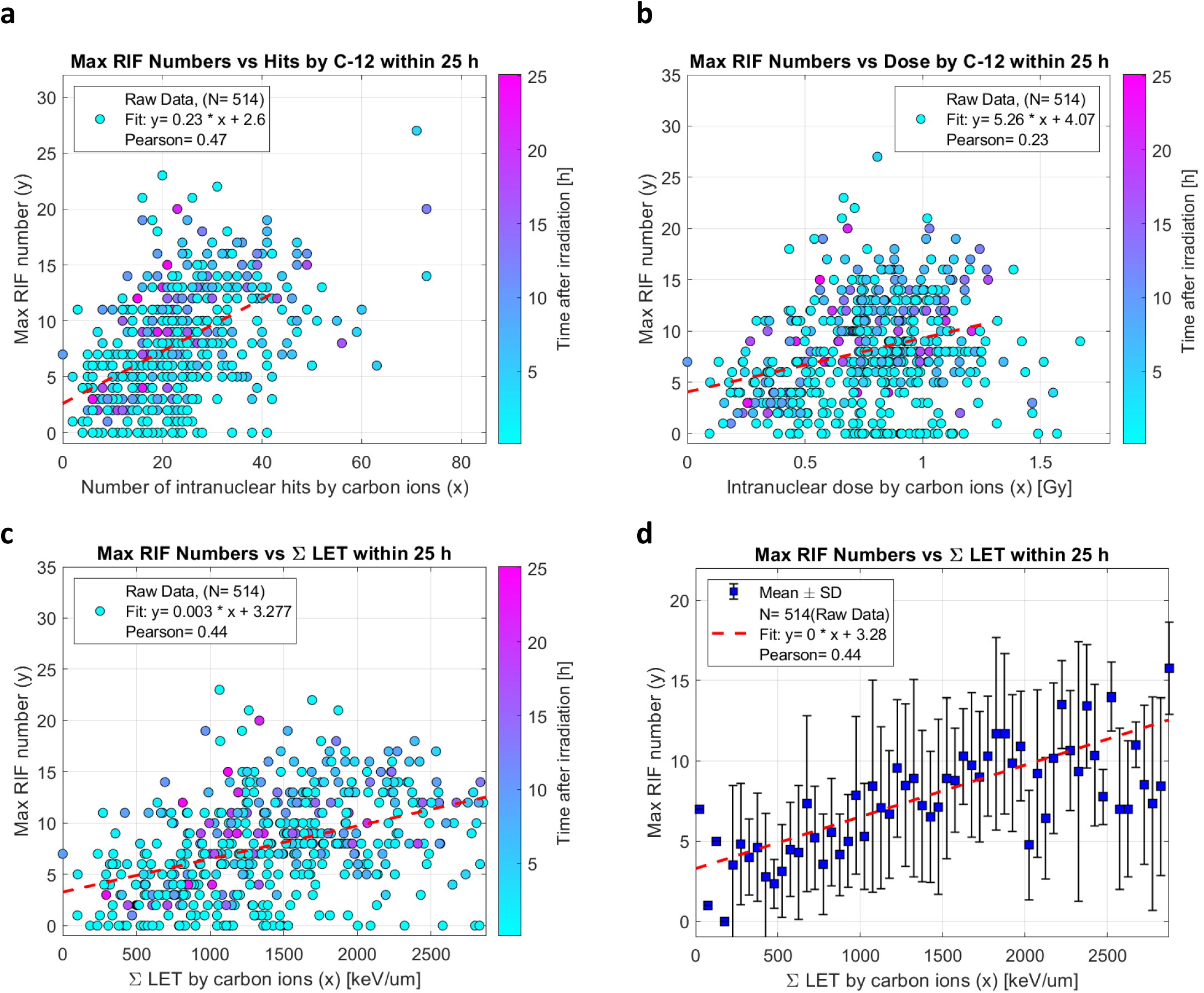
*Cell-Fit-HD*^*4D*^ enables correlation of physical ion beam parameters with molecular dynamics of individual tumor cells. **(a)-(c)** Raw data for the correlation of hits/dose and ΣLET with maximum RIF numbers within 25 h post irradiation and linear regression (red dashed line). The first time point of the occurrence of maximum RIF number in a cell nucleus is encoded by the colour bar. **(d)** Mean values and SD for the correlation of ΣLET with maximum RIF numbers. For the computation of the Pearson-r correlation and linear regression analysis the raw data was used. Here, the hit number, dose and ΣLET were limited to 42 hits, 1.3 Gy and 2875 keV/um, respectively to ensure sufficient number of data points. Only mother cells were considered in the data analysis.

**Figure SI8.**
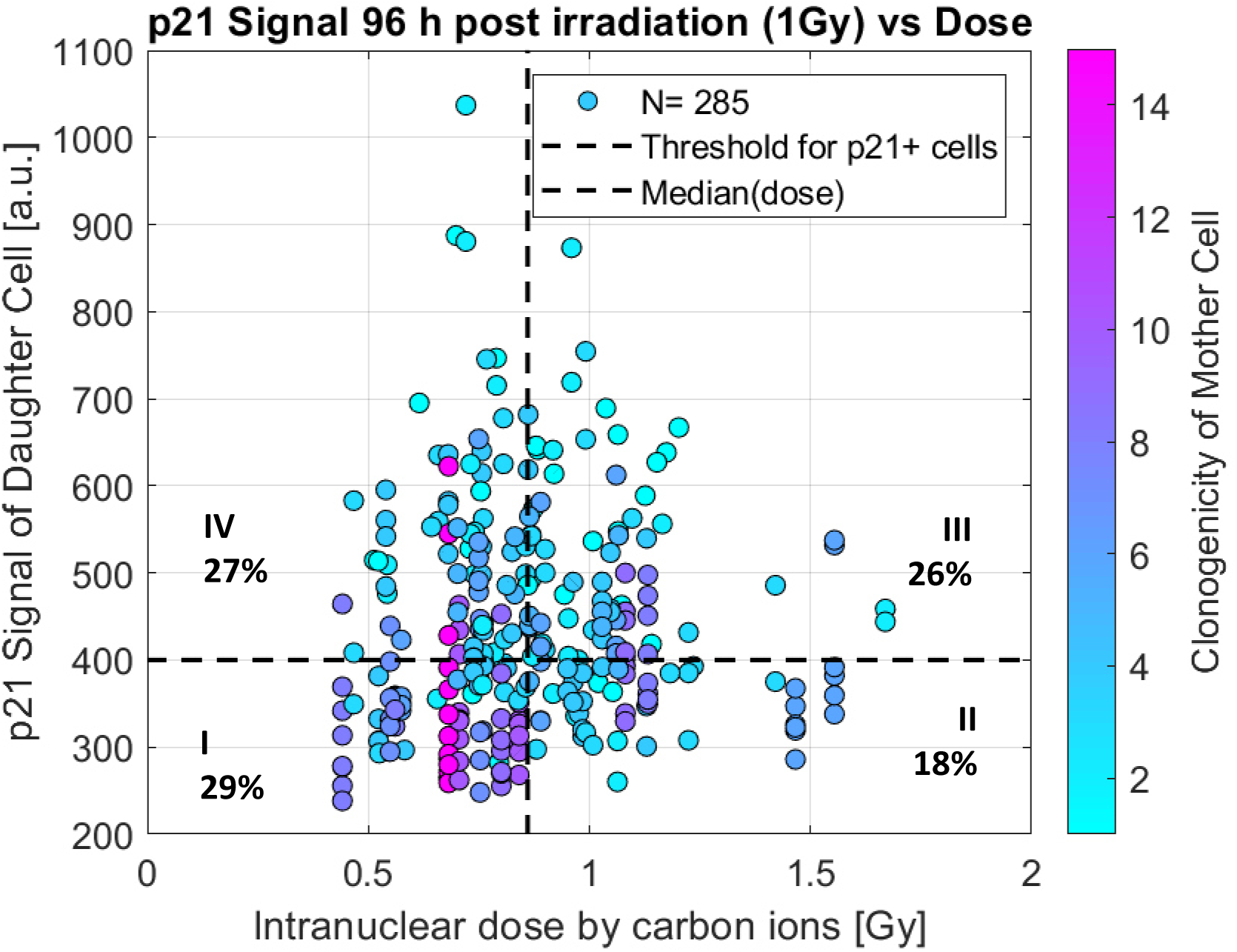
*Cell-Fit-HD*^*4D*^ combines single-cell dosimetry with molecular and cellular dynamics of individual tumor cells. Multi-dimensional correlations of physical beam (dose)- and molecular (p21 status 96 h after irradiation) parameters to predict proliferation potential and to find potential marker for radiation-resistance. Clonogenicity: number of progeny cells deriving from a single mother cell. The plot was divided into four quadrants (I-IV): median dose= 0.82 Gy, p21 threshold= 400 a.u. The portion of the total population is given for each quadrant.

**Figure SI9.**
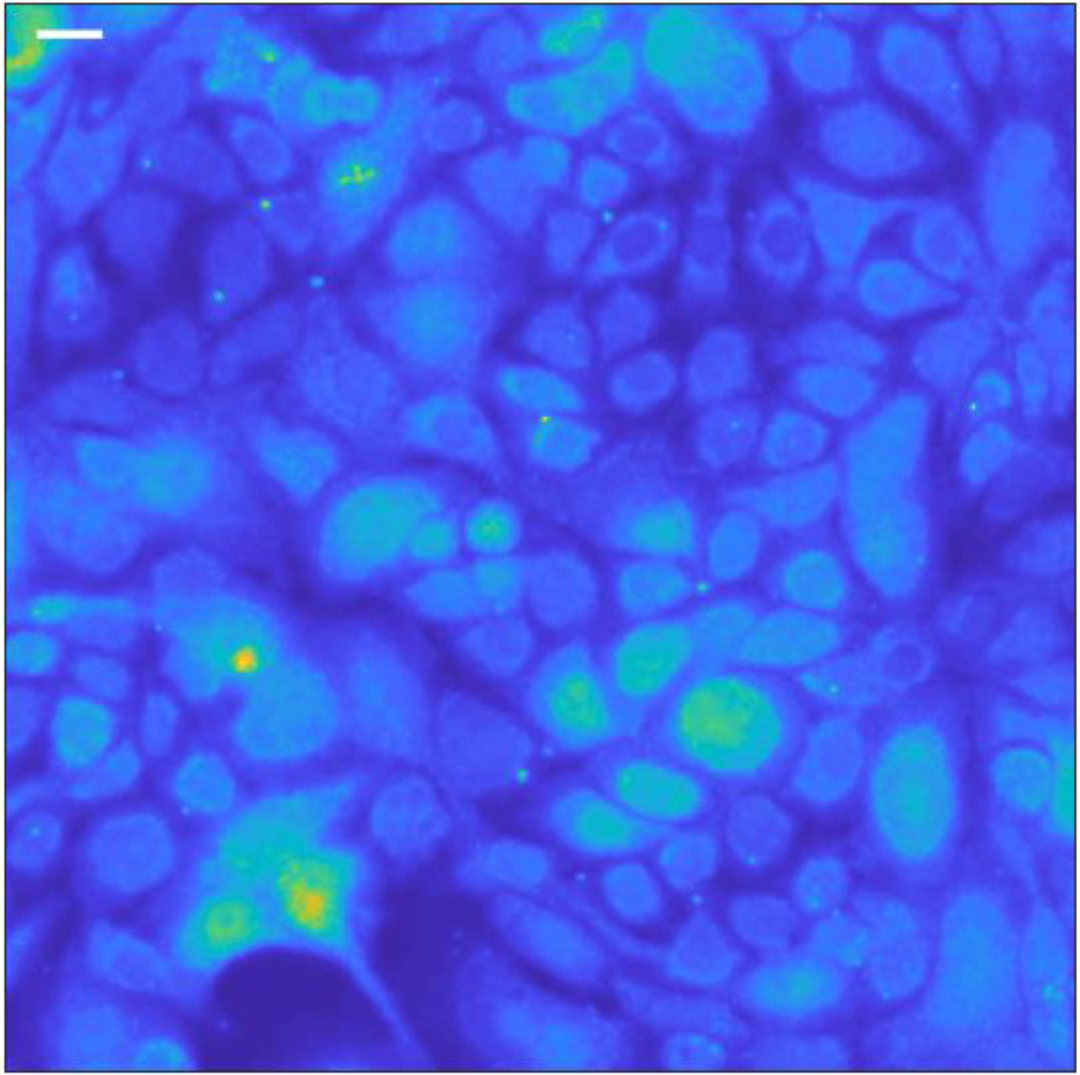
P21 Signal 96h after irradiation (1Gy SOBP, carbon ion). Exemplary section of A549 cell layer with fluorescent p21 signal. Scale bar, 20 µm.

