## Supplementary Information for "*The biomedical sensor Cell-Fit-HD^4D^*, reveals individual tumor cell fate in response to microscopic ion deposition"

We performed a series of tests in order to validate both sensor architectures and the corresponding analysis routines.

#### Alternative design of *Cel-Fit-HD$^{4D}$*: Mounting-architecture

##### ***Rationale behind mounting architecture***

The main rationale behind developing this architecture was to account for the limitation in the wavelength spectrum of potential fluorescent dyes in the cell layer. A recording of fluorescent dyes in the cell layer by widefield microscopy with excitation wavelength close to 405 nm would considerably alter the signal of the track spots in the wafer. An excitation wavelength close to 620 nm would also principally interfere with the emission signal of the wafer (*Paragraph ion track reconstruction $Al_2O_3$:C,Mg based FNTD*) at the border region between wafer and cell layer. We therefore aimed for architecture to separate the physical (FNTD) and the biological (cell layer) compartment. By introducing sufficient distance (≥ 25 µm) between both compartments, the interference of both read-outs could be dramatically reduced. Replacing the wafer with a glass bottom for cell culturing resulted in better SNR.

##### ***Corresponding design of Mounting-architecture***

The design of the Mounting-architecture is illustrated in Figure SI3a. The rationale behind this construction was to minimize potential displacement between the now separated cell layer and the FNTD during irradiation. This displacement gets directly reflected in the uncertainty in ion track reconstruction in the cell layer. A single well of 12-well-glass bottom dish (MatTek) was covered with a lid (framed in red in Figure SI3a). It contains a stamp with the FNTD. The lid contains small apertures to fill the insert of the well with culture medium (indicated in magenta). Inert and non-absorbing material (glass and alumina) and little number of adhesive joints were used in this lid. To ensure optimal cell physiological conditions during live-cell imaging the lid (including the FNTD) gets removed after the initial read-out (step 1 in Figure 3a, *Paragraph Workflow of Cell-Fit-HD$^{4D}$*). For the removal, force is applied gently at the adhesive joints (labelled in yellow in Figure 4a) by a pair of tweezers.

Different designs made of plastic and manufactured by 3D printing were initially tested. Optimal culturing conditions for the cell layer were achieved. Before use we placed the sterilized architecture in culture medium at humidified atmosphere for approximately 48h. Potential displacement of several µm between cell layer and FNTD during the time interval between irradiation and initial read-out caused by thermal expansion (small temperature gradient during irradiation) and absorption of the medium by the plastic was still possible.

##### ***Sandwich construction for validation of Mounting-architecture***

The ions are principally travelling on straight lines when traversing the biomedical sensor. We still performed a validation experiment to prove the accuracy in ion track reconstruction using the read-out signals of the FNTD in the Mounting-architecture (Figure SI3a). For this purpose, we substituted the glass bottom in the Mounting-architecture with a FNTD in the form of a thin wafer. For this step, we used histo-acryl glue (8 ul, *paragraph Design of Cell-Fit-HD$^{4D}$ and manufacturing approaches,* Figure SI3b). The distance between FNTD and wafer (Δh) was approximately 45 µm. We performed carbon ion irradiation using identical irradiation plan (paragraph *Irradiation*) but without stopping material (PMMA). We reconstructed the ion trajectories both in the wafer (blue arrows) and in the FNTD (red arrows) using the ion track reconstruction routines described in *Ion track reconstruction and dose calculation in cell nucleus*. Additionally we extrapolated the ion trajectories from the FNTD onto the wafer (red dashed lines) surface and tested the spatial overlap of their coordinates with the coordinates of the ion trajectories reconstructed in the wafer.

Since we were using the identical read-out protocol as for live-cell imaging (*Paragraph Workflow of Cell-Fit-HD$^{4D}$*) we performed initial read-out of the wafer and of the FNTD by widefield microscopy within a single step. The Mounting-architecture was then disassembled and the wafer and FNTD were scanned independently by CLSM. A tile scan and subsequent stitching routine was performed only in the confocal read-out of the wafer. Both recorded confocal image stacks were registered with the data set recorded by widefield microscopy in the initial read-out. The associated ion trajectories (originating from the FNTD and from the wafer) were identified and linked using the nearest neighbour algorithm in *Matlab*. This pairing was validated manually. We tested spatial overlap of the reconstructed ion trajectories within two regions, resulting in the mean displacement of 1.02 µm and 1.30 µm, respectively. We, in general, overestimated this error since we performed image registration twice (instead of once, *Paragraph Workflow of Cell-Fit-HD$^{4D}$*). In addition, we also neglected possible expansion of the glue (*paragraph Thermal expansion of the glue*) and we were conducting the experiments at RT (room temperature).

Due to the industrial manufacturing of the glass bottom multiwell dishes (MaTek) with limited precision, the distance Δh between glass bottom and FNTD varies. This, in principle, does not disturb the ion track reconstruction, but is reflected in greater prediction intervals (PIs) [16].

#### *Thermal expansion of the glue*

We investigated the influence of the viscosity of the glue used in assembling of the Mounting-architecture on spatial correlation of the information of the biological and physical compartment. In particular, we investigated, whether dilatation of the glue (used to mount the FNTD) based on temperature change during the irradiation of *Cell-Fit-HD$^{4D}$* could influence the ion track reconstruction in the cell layer. For the validation experiment the glass bottom in the Mounting-architecture was replaced by a wafer. We principally tested histo-acryl glue (B. Braun GmbH, Cat-No. 9381104) and *Sekundenkleber Blitzschnell Präzision* (UHU).

We mimicked the assembling, irradiation and read-out procedure of *Cell-Fit-HD$^{4D}$*: We assembled the Mounting-architecture and incubated the multiwell plate filled with culture medium (without cell layer) in the incubator (humified atmosphere) for approximately 24 h (step 1). After replacing the culture medium and sealing the multiwell plate with Parafilm under RT (step 2) we stored the plate again in the incubator for 6 min (step 3). We simulated the irradiation by placing the biomedical sensor in an upright position for 6 min under RT (step 4). We stored the biomedical sensor for additional 2 min at RT to account for the transfer from the irradiation facility to the microscope (step 5). In the last step, we ran the initial read-out of the biomedical sensor at 37°C (step 5). At the end of steps 2-5, we performed imaging of the *spinels* located in the wafer and in the FNTD by widefield microscopy. For steps 2-5 we defined five control point pairs (*cpps*) using the recorded *spinels* in the wafer and in the FNTD. We calculated the distance in the horizontal plane for each *cpp* for each image acquisition to detect any spatial shift during the temperature change caused by the changing viscosity of the glue. We identified the shift in the critical time span between irradiation and initial read-out to be less than 1 µm.

#### *Signal-to-Noise ratio (SNR)*

We investigated the signal-to-noise ratio (SNR) in the live-cell imaging using the Mounting-architecture and the wafer. The SNR in both designs is sufficient for detection of molecular and cellular kinetics labelled by fluorescent dyes including RIF formation, cell migration and cell division. There is a great inter-cellular variation in the signal of the recorded cell nuclei. This is based on the substantial variation of 53BP1 expression within the cell population. The cell nuclei cultured on the wafer principally appeared to be dimmer in the imaging data compared to the nuclei cultured on the glass bottom in the Mounting-architecture. The edges (gradient) of the foreground object in the Mounting-architecture appeared to be steeper compared to the wafer architecture. The SNR gained in both designs was however sufficient to perform nucleus segmentation and cell tracking using the *Matlab* routine *LSetCellTracker* and RIF detection using *Trainable Weka segmentation* plugin for *ImageJ [47]*. The ratio of mean number of recorded RIFs in a cell nucleus at 0.87 h after irradiation using the Mounting-architecture and the wafer architecture was ~ 4.39/3.24 = 1.35. This ratio arises from small and dim RIFs in the wafer architecture which (up to now) cannot always be detected by the *Trainable Weka segmentation* plugin. The large RIFs – presenting the relevant damage – are, however, visible in both designs. The underlying RIF dynamics is still very similar in both architectures.

It was not always possible for the segmentation routine (*LSetCellTracker*) to identify the correct edges of the nuclei. The resulting areas sometimes appear too large or small. Yet, this observation is rather subjective as there is no explicit definition of the "correct" edges of the nuclei. We, therefore, used identical segmentation parameters for the experiments using identical architectures (Mounting-architecture or wafer) to get consistent and comparable results. Moreover, under- and over-segmentation of the cell nuclei in both architectures are balanced.

### Cell viability

We investigated, whether the cell viability on the wafer (Figure 2) differs from the cell viability using the glass bottom (Mounting architecture, Figure SI3a). Cell number dynamics was recorded for each imaging field (Figure SI4a). The area under the curve was thus computed and normalized by the number of cells in the initial time frame (Figure SI4b). For this purpose, a binary mask was created by *global thresholding* using the recorded fluorescent signal of 53BP1 and modified by the functions *dilation* and *erosion* in *Matlab*. To separate individual cell nuclei watershed segmentation was used as presented in [48]. This algorithm turned out to be more sensitive in detection of nuclei with low 53BP1 expression compared to *LSetCellTracker* .

Our findings agree with former studies published in [25]. The proliferation curves and boxplots reflect the principal heterogeneity in cell proliferation dynamics. For statistical testing the number of imaging fields could be increased.

### RIF frequency

We investigated, whether the extracted RIF (53BP1) dynamics gained from the read-out signal of wafer and the Mounting-architecture of *Cell-Fit-HD*$^{4D}$ was similar. With this test, we were looking for hints indicating different molecular dynamics since the cells were cultured on different substrates (glass vs alumina). As depicted in Figure SI1, cells cultured on alumina (wafer) or glass (Mounting-architecture) and being exposed to 0.5 Gy (C-12) showed similar RIF dynamics. The mean number of RIFs in a nucleus peaked at approximately 50 min post irradiation, followed by a decline. Cells cultured on wafer and being exposed to 1Gy (C-12) exhibit principally similar RIF dynamics with a delayed maximum at approximately 67 minutes after irradiation. The maximum mean RIF numbers are shifted towards greater values by a factor of approximately 2.01 when comparing the data set from 0.5Gy and 1 Gy wafers. The shift is - as expected - caused by the higher dose deposition in the 1 Gy experiment. The difference in peak values by a factor of approximately 1.4 when comparing the 0.5 Gy wafer and 0.5 Gy Mounting-architecture datasets is most likely caused by the decreased SNR in the imaging data of the 0.5 Gy wafer experiment. To principally account for the difference in SNR, the *Trainable Weka Segmentation* Tool was trained for the wafer and the Mounting-architecture datasets individually.

### Tracking using *LSetCellTracker*

For nucleus segmentation and cell tracking we used the *Matlab* routine *LSetCellTracker* (*paragraph LSetCellTracker in Materials and Methods*). We performed automated cell-tracking of the imaging data recorded by live-cell imaging in a time interval of approximately 96 h. To assess the quality of tracking we were manually screening for errors in the generated cell division trees. Within this scope, the daughter cells in the last time frame were assigned to the corresponding mother cells (in total 531 mother cells) in the initial time frame of all imaging data acquired using the wafer architecture. Approximately 85% of daughter cells were assigned to the corresponding mother cells (including no error in cell division). Approximately 15 % of daughter cells were falsely assigned. Approximately 7 % of this population were at the transition between correct and miss-assignment.

The variation of 53BP1 expression within the cell population and a close contact of neighbouring cells were identified to be the two main sources of errors. In the latter case neighbouring cells were morphologically identified as a single cell. These sources of errors are common issues in cell tracking and are not directly related to the architecture of our biomedical sensor. Optimization of cell tracking could be achieved by using fluorescent 53BP1 constructs with homogenous expression and by applying artificial intelligence or neural network approaches.

To minimize the cell tracking errors, we applied non-supervised and automated cell tracking until 25 h after irradiation.

### γ-H2AX vs endogens 53BP1

For the development of the architecture of *Cell-Fit-HD*$^{4D}$ we used the 53BP1 fluorescent damage protein accumulating at DNA damage sites induced by the ion irradiation. 53BP1 is recruited to DSB sites in a slightly delayed fashion compared to γ-H2AX [29]. We determined the spatial overlap of the 53BP1- with γ-H2AX signal. In the first step, we determined that 73.9% of the γ-H2AX foci correlate well with 53BP1 foci, both gained by immunofluorescent labelling and confocal microscopy. This discrepancy is most likely explained by high-resolution-dependent detection of spontaneous, short-living γH2AX-foci caused by cell-inherent processes like replication stress.

Expression of 53BP1-mCherry can vary substantially among cells. However, since foci detection occurred in a signal accumulation-dependent manner and with the help of machine learning, which is mainly independent of total intensity, no limitations were expected on that account. Notably, heavy ion irradiation produces complex DSB clusters which directly lead to formation of large, high-intensity RIF [49] what minimizes the possibility for missed RIFs. Nonetheless, 53BP1-mCherry-signals were aligned with immunofluorescently stained 53BP1-protein. We determined that 98.8% of large 53BP1 RIF (area > 8 pixel) labeled by immunofluorescence overlap with the 53BP1-mCherry foci signal. Here, the identical widefield microscope as for the read-out of the biomedical sensor *Cell-Fit-HD$^{4D}$* was used (*Paragraph Workflow of Cell-Fit-HD$^{4D}$*).

### Precision of microscopy stage movement

We validated the precision of the microscopy stage movement since it has a major impact on the *Workflow of Cell-Fit-HD$^{4D}$* (*Paragraph Workflow of Cell-Fit-HD$^{4D}$*). An identical imaging position was recorded over time in the brightfield mode by widefield microscopy (IX83, Olympus) using the wafer-architecture A global coordinate system was defined by the *spinels*. Stage movements during sequential imaging of identical positions were computed in a time interval of approximately 72 h. The norm vector of the *spinel* positions (detecting intensity-weighted centroids of the *spinels*) in the maximum intensity projection between two consecutive time points was computed. The norm vector between the initial and second time point was greater than 1 µm. This parameter decreased rapidly at later time points. The dimension (>> 1 µm) of the moving vector at early time points reflects the needs for the recording of the initial stack and consecutive image registration to spatially correlate the ion traversals with cellular information (*Paragraph Workflow of Cell-Fit-HD$^{4D}$*). Neglecting this movement (whether by stage uncertainty or thermal movement, expansion) would result in a degradation of the accuracy in computation of the spatial energy deposition by the ions in the cell layer. The recording of the initial stack of relatively long acquisition time (and hence improved accuracy) is principally in conflict with the acquisition of high number of imaging fields in a short time interval after irradiation. A short time interval of approximately 5 minutes is necessary to minimize further sources of errors (e.g. cell migration) compromising the spatial correlation of ion traversals with individual cell response.

### Assessment of error sources/ uncertainties in spatial correlation

Below, we listed all potential sources of errors which principally affect the determination of the correct position of the ion traversal in the cell layer and hence the number of intranuclear hits and computed dose:

- Segmentation error $\Delta s_i$ of the cell nuclei
- Cell migration $\Delta d_i$ on the wafer in the time interval between irradiation and initial read-out
- Error $\Delta r_i$in registration procedure (including stitching of single imaging tiles),
- Uncertainty $\Delta z_i$ of the correct vertical position of the cell layer,
- Uncertainty in track reconstruction expressed by the prediction interval $\mathrm{PI}_{x,y}$.

Migration ($\Delta d_i$) and the segmentation errors ($\Delta s_i$) are based on representative numbers of cell nuclei: $N = 183$ and $N = 202$, respectively.

We developed a Monte Carlo-based simulation to estimate the total error in the computation of the number of intranuclear hits. The underlying algorithm was published in [50]. The simulation was implemented in *Matlab*. To determine the error distribution for each individual cell, the simulation was looped with $N = 1000$ for each cell. The algorithm for individual cell nuclei consists of the following steps:

1. Load binary mask $BI_i$ (created by the *LSetCellTracker*) of the maximum intensity projection $A_i$ of nucleus$_i$ in the initial frame after irradiation.
2. Choose a migration distance $\Delta d_i$ by randomly sampling from a list with migration distances occurring within a time interval of 35 minutes. Set migration direction by randomly sampling a polar angle $\phi_i$. Shift all foreground pixels by the corresponding translation vector (rounded to integer values).
3. Choose a segmentation error $\Delta s_i$ of the nucleus by randomly sampling from list with segmentation errors. Depending on the sampling value (over-segmentation, under-segmentation, and correct segmentation) the binary mask $BI_i$ is enlarged or reduced.
4. Define vertical position $\hat{z}$ of the cell layer (maximum intensity projection) with respect to the wafer surface by randomly sampling the corresponding error $\Delta z_i$. Select the corresponding track positions and prediction intervals $\mathrm{PI}_x$ and $\mathrm{PI}_y$ in $\hat{z}$.
5. Recalculate $\mathrm{PI}_x$ and $\mathrm{PI}_y$ in corresponding normal distribution with $\mathrm{PI}_{x/y} = 2\ \mathrm{SD}_{x/y}$. Compute $\widehat{PI_x}$, $\widehat{PI_y}$ by sampling from the corresponding normal distribution.
6. Choose registration error $\Delta r_i$ (rounded to integer values) by randomly sampling from list with the Euclidean distances between the control point pairs. Set translation direction by randomly sampling a polar angle $\phi_i$.
7. Determine all actual track positions $(x, y)$ by translating the track positions in $\hat{z}$ with $\widehat{PI_x}$, $\widehat{PI_y}$, and $\Delta r_i$.
8. Determine number of intersections of track positions $(x, y)$ with the modified binary mask $BI_i$.

At the end of the simulation the mean number and standard deviation of the distribution of intranuclear hits for each cell nucleus were computed.

The mean total error is much smaller than the dimensions of cell nuclei (having the diameter of 10 µm).
